# Profiling chromatin accessibility in formalin-fixed paraffin-embedded samples

**DOI:** 10.1101/2021.07.08.450856

**Authors:** Vamsi Krishna Polavarapu, Pengwei Xing, Hua Zhang, Miao Zhao, Lucy Mathot, Linxuan Zhao, Gabriela Rosen, Fredrik J. Swartling, Tobias Sjöblom, Xingqi Chen

**Author notes:** Contributed equally (Vamsi Krishna Polavarapu, Pengwei Xing, Hua Zhang).

## Abstract

Archived formalin-fixed paraffin-embedded (FFPE) samples are the global standard format for preservation of the majority of biopsies in both basic research and translational cancer studies, and profiling chromatin accessibility in the archived FFPE tissues is fundamental to understanding gene regulation. Accurate mapping of chromatin accessibility from FFPE specimens is challenging because of the high degree of DNA damage. Here, we first showed that standard ATAC-seq can be applied to purified FFPE nuclei but yields lower library complexity and a smaller proportion of long DNA fragments. We then present FFPE-ATAC, the first highly sensitive method for decoding chromatin accessibility in FFPE tissues that combines Tn5-mediated transposition and T7 *in vitro* transcription. The FFPE-ATAC generates high-quality chromatin accessibility profiles with 500 nuclei from a single FFPE tissue section, enables the dissection of chromatin profiles from the regions of interest with the aid of hematoxylin and eosin (H&E) staining, and reveals disease-associated chromatin regulation from the human colorectal cancer FFPE tissue archived for more than 10 years. In summary, the approach allows decoding of the chromatin states that regulate gene expression in archival FFPE tissues, thereby permitting investigators, to better understand epigenetic regulation in cancer and precision medicine.

## INTRODUCTION

Decoding the landscapes of chromatin regulatory elements in human disease, specifically cancer, is of critical importance in preclinical diagnosis and treatment(Qu et al. 2017). Recently developed technologies, such as the assay for transposase-accessible chromatin by sequencing (ATAC-seq) (Buenrostro et al. 2013)and DNase I hypersensitivity sequencing (DNase-seq)(Jin et al. 2015), allow profiling of chromatin accessibility in cells and frozen tissues. Archived formalin-fixed paraffin-embedded (FFPE) tissues are the global standard format for preservation of the majority of biopsies in basic research and translational cancer studies(Fox CH 1985), and it has been reported that more than 20 million FFPE specimens are newly archived every year in the United States alone(Waldron et al. 2012). Accordingly, profiling gene regulation in the archived FFPE tissue can be invaluable for translational cancer research. Chromatin structure is still preserved during FFPE sample preparation and long-term storage(Fanelli et al. 2010; Jin et al. 2015; Cejas et al. 2016). However, it has proven difficult to apply the currently available highly sensitive chromatin accessibility decoding technologies to FFPE tissue samples because of the high degree of DNA damage that occurs during sequencing library preparation of these samples (Chin et al. 2020). Moreover, it is desirable that a minimum number of FFPE tissue sections be used in the analysis, as the tissues of interest are limited. The currently required input for chromatin structure studies from FFPE samples is either couples of tissue sections or whole tissue block (Fanelli et al. 2010; Jin et al. 2015; Cejas et al. 2016), and this precludes conducting analyses at high resolution. To this end, we developed FFPE-ATAC, the first highly sensitive method for decoding the chromatin accessibility in FFPE tissues, by combining the Tn5-mediated transposition and T7 *in vitro* transcription.

## RESULTS

### Standard ATAC-seq on FFPE samples

During formalin fixation, the formaldehyde in the formalin reacts with primary amines to form Schiff bases, and with the amides to form hydroxymethyl compounds, resulting in the formation of large chromatin complexes(Fox CH 1985). To decode the chromatin states in the FFPE samples, it is essential to disrupt these chromatin complexes using reverse cross-linking(Fanelli et al. 2010; Cejas et al. 2016). In standard ATAC-seq for live cells or frozen tissues, accessible genomic sites are amplified and enriched through the polymerase chain reaction (PCR) by using primers that hybridize with the universal Tn5 adaptors(Buenrostro et al. 2013). In our previously established ATAC-see (Chen et al. 2016) and Pi-ATAC(Chen et al. 2018) technologies, we used a reverse cross-linking step to remove mild formaldehyde cross-linking and performed ATAC-seq in the mildly fixed cells at the bulk and single-cell levels. However, we learned that the reverse cross-linking step can cause a high degree of DNA damage and introduce DNA breaks in extensively fixed cells and the FFPE tissues (**Fig. 1A**) (Martelotto et al. 2017). Furthermore, we assumed that if such DNA breaks occur at accessible chromatin sites in FFPE tissues, and, if so, that this might hamper PCR amplification of those accessible chromatin sites with the standard ATAC library preparation strategy (**Fig. 1A**). To test our hypothesis, we developed an optimized protocol for the isolation of high-quality nuclei from mouse liver and kidney FFPE tissue sections with 20 µm in thickness (**Supplemental Fig. S1A, S1B**, see **Methods**). Following the reverse cross-linking strategy, we indeed observed many DNA breaks in the genomic DNA purified from isolated FFPE nuclei (**Supplemental Fig. S1C**). In addition, only short fragments were obtained when the standard ATAC-seq procedure was used on 50 000 nuclei isolated from mouse FFPE liver and kidney tissues (**Supplemental Fig. S1D, S1E**; see **Methods**), suggesting that DNA breaks indeed occur at accessible chromatin sites with long DNA lengths and that this further hampers PCR amplification of those regions (**Fig. 1A**). We then sequenced the libraries obtained through standard ATAC-seq on isolated FFPE nuclei (**Supplemental Fig. S2A**), and prepared standard ATAC-seq libraries on frozen samples collected from the same mouse liver and kidney samples as FFPE samples (**Supplemental Fig. S2B**). Then, we compared the sequencing libraries obtained by standard ATAC-seq on FFPE samples with those obtained by standard ATAC-seq on frozen samples (**Fig. 1B-I, Supplemental Fig. S2, Supplemental Fig. S3, Supplemental Code**). This resulted in several findings. **i**) The proportion of long DNA fragments (longer than 146 bp) obtained from standard ATAC-seq on FFPE samples (30.76% +/- 1.38% for liver and 43.15% +/- 0.5% for kidney) was lower than that obtained from standard ATAC-seq on frozen samples (50.02% +/- 4.7% for liver and 59.17% +/- 2.57% for kidney) (**Fig. 1B, 1C**). Furthermore, the proportion of mononucleosome fragments enriched at transcription start sites (TSS) was also lower from the standard ATAC-seq on FFPE samples (**Supplemental Fig. S2C, S2D**). **ii**) The library complexity obtained from standard ATAC-seq on FFPE samples was much lower than that obtained through standard ATAC on frozen samples (**Fig. 1D, 1E**). **iii**), The proportion of mitochondrial reads obtained from standard ATAC-seq on FFPE samples (27-42%) was much higher than that obtained through standard ATAC-seq on frozen samples (2-6%) (**Fig. 1F, 1G**). Since all of the ATAC-seq libraries were prepared from purified nuclei, the sequencing libraries should contain very limited amounts of mitochondrial DNA. The high proportion of mitochondrial reads obtained through standard ATAC-seq on FFPE samples may be due to the fact that library complexity from genomic DNA in FFPE samples is low, and PCR amplification enriches a high percentage of mitochondria. **iv**), High TSS enrichment scores (score number: 27-30) and high number fraction of reads in peaks (FRiP) (over 40%) were obtained from standard ATAC-seq on FFPE samples (**Fig. 1F, 1G, Supplemental Fig. S2E, S2F**). Standard ATAC-seq on FFPE samples also showed good genome-wide correlation with the results of standard ATAC-seq on frozen tissue (mouse liver: *R* = 0.87, mouse kidney: *R* = 0.85, **Fig. 1H, 1I**), and the distribution of sequencing reads in the genome from standard ATAC-seq on FFPE samples was similar to the distribution obtained by standard ATAC-seq on frozen tissue (**Supplemental Fig. S2G, S2H**). In addition, a large proportion of the peaks obtained by standard ATAC-seq by standard ATAC-seq on FFPE samples and standard ATAC-seq on frozen tissue overlapped (**Supplemental Fig. S3A, S3B**). Exclusive peaks from standard ATAC-seq on FFPE samples and standard ATAC-seq on frozen tissue are distributed randomly in the genome and display similar enrichments of transcription factors (**Supplemental Fig. S3A, S3B**). **v**) However, we noticed that a proportion of the accessible regions are much more open in frozen samples than in FFPE samples (**Fig. 1H, 1I**). On differential peak analysis (Log_2_ (fold change) > 3, p < 0.01) (Supplemental Code)(Love et al. 2014), many more accessible chromatin regions were identified in the frozen samples (n = 1598 in mouse liver and n = 495 in mouse kidney), but almost no more accessible chromatin regions were identified in the FFPE samples (n = 0 in mouse liver and n = 3 in mouse kidney) (**Fig. 1H, 1I**), suggesting that standard ATAC-seq on FFPE samples failed to detect a proportion of the accessible chromatin sites. To further investigate whether the more accessible regions in standard ATAC-seq on frozen samples represent sites at which DNA breaks occurred in the FFPE samples, we calculated the number of sequencing reads obtained for those regions in standard ATAC-seq on FFPE samples and found that for 66.33% (1060/1598) of those regions in FFPE mouse liver and 55.77% (256/459) of those regions in FFPE mouse kidney, no sequencing reads were detected (**Supplemental Table S1, Supplemental Table S2**). This strongly suggests that DNA breaks potentially occur at those sites in FFPE samples and further hamper PCR amplifications of them. We also noticed that the more accessible regions in standard ATAC-seq on frozen samples were mainly located at regions distal (>10 kb) to the TSS (**Fig. 1H, 1I**).

**Figure 1:**
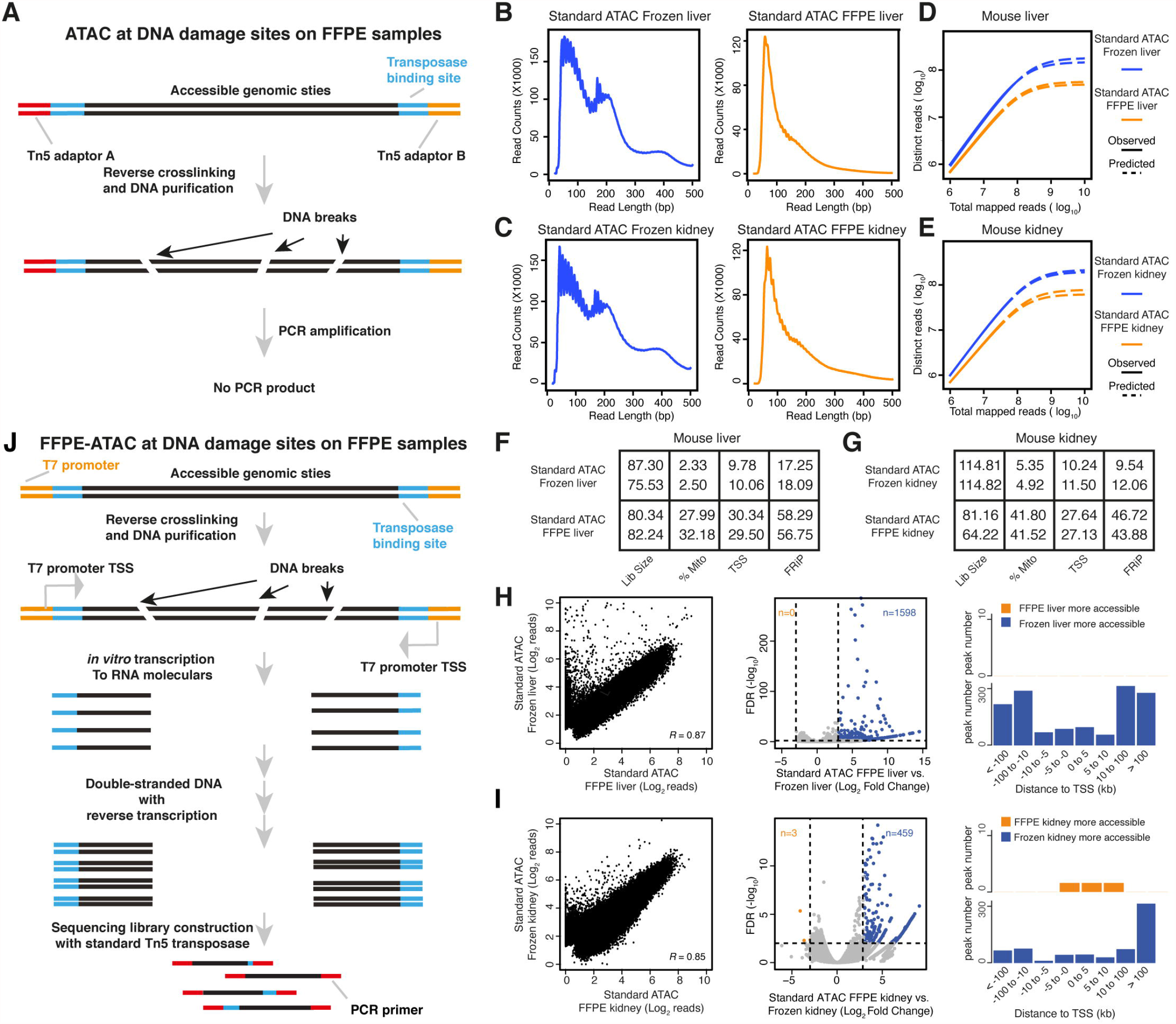
Standard ATAC-seq on FFPE samples and design of FFPE-ATAC. (**A**), DNA damage on accessible chromatin sites in FFPE samples hampers PCR amplification in standard ATAC-seq on FFPE samples. (**B-E**), Comparison of DNA fragment size distribution (**B, C**) and library complexity (**D, E**) from standard ATAC-seq on frozen mouse liver and kidney, and standard ATAC-seq on FFPE mouse liver and kidney. (**F, G**), Quality control metrics of standard ATAC-seq on frozen mouse liver (**F**) and kidney (**G**), and standard ATAC-seq on FFPE mouse liver (**F**) and kidney (**G**). Lib size = total sequencing reads of sequencing library (million); %Mito = percentage of mitochondria; TSS = enrichment score at transcription start sites (TSS); FRiP = fraction of reads in peaks. (**H, I**), Comparison of chromatin accessibility between standard ATAC-seq on frozen samples and FFPE samples. Left: Genome-wide comparison of accessible chromatin regions. *R* = Pearson’s correlation. Middle: Differential peak analysis between standard ATAC-seq on frozen samples and FFPE samples. FDR = false discovery rate. Right: Distribution of the more accessible regions from frozen and FFPE mouse samples across transcription start sites (TSS). (**J**), Design of FFPE-ATAC by combining T7-Tn5 transposase tagmentation and T7 *in vitro* transcription.

Taken together, our results show that the transposase-mediated technology, ATAC-seq, can be applied to FFPE samples consisting of nuclei isolated through an optimized procedure. However, we learned that DNA breaks at accessible chromatin sites in FFPE samples potentially hamper PCR amplification of these regions when standard ATAC-seq is used. We concluded that standard ATAC-seq libraries on FFPE samples have lower library complexity and a lower proportion of long DNA fragments, and lack a proportion of the accessible chromatin sites compared with libraries prepared by standard ATAC-seq on frozen samples.

### The design of FFPE-ATAC

To increase the library complexity and rescue lost accessible regions in standard ATAC-seq on FFPE samples, we developed FFPE-ATAC to decode chromatin accessibility in FFPE tissues by combining Tn5-mediated transposition and T7 *in vitro* transcription (IVT) (**Fig. 1J**). During Tn5 transposition in FFPE samples, Tn5 adaptors are inserted into the genome after FFPE sample preparation; they are therefore unlikely to undergo the DNA breakage that occurs during reverse cross-linking of FFPE samples and should therefore remain at the ends of broken accessible chromatin sites after reverse cross-linking. We reasoned that by adding a T7 promoter sequence to the Tn5 adaptor (**Fig. 1J, Supplemental Fig. S4A**) we could use IVT to convert the two ends of the broken DNA fragments to RNA molecules before preparing sequencing libraries from the IVT RNAs, and further decode the Tn5 adaptors’ insertion sites in the genome (**Fig. 1J**). Through this strategy, we could decode the flanking sequences of the accessible chromatin despite the fact that there were breaks between adjacent pairs of T7-T5 adaptor insertion sites. It was found that Tn5 activity is very robust, given the different sequence modifications on the Tn5 adaptor(Chen et al. 2016; Sos et al. 2016; Chen et al. 2017; Xie et al. 2020; Payne et al. 2021). Thus, we designed, produced, and optimized a Tn5 adaptor with an added T7 promoter sequence, termed T7-Tn5 (**Supplemental Fig. S4A**, see **Methods**). T7-Tn5 retains the activity of the standard Tn5 (**Supplemental Fig. S4B**, see **Methods**). To test our hypothesis that the T7-Tn5 adaptors remain at the ends of the accessible chromatin DNA fragments despite the DNA breaks that result from reverse cross-linking, we performed IVT on single nuclei obtained from FFPE samples of mouse liver and kidney after T7-Tn5 transposition. We found that RNA fractions from these two FFPE tissues contained both short and long RNA (**Supplemental Fig. S4C**). This result suggests that the T7 promoter is still present at the ends of the broken accessible chromatin sites in the long-term fixed FFPE samples after reverse cross-linking and that the insertion sites of T7-Tn5 adaptors in the genome could be decoded in RNA molecules from IVT even when only one T7-Tn5 adaptor was present at the end of the broken DNA molecules. Our results indicate that use of a combination of Tn5 transposition and T7 IVT could be of value for performing FFPE-ATAC and that it potentially rescues broken DNA fragments in FFPE samples at accessible chromatin regions.

### Proof of concept of FFPE-ATAC with mouse FFPE liver and kidney samples

Next, we proved the principle of FFPE-ATAC using sets of 500-50 000 nuclei purified from individual FFPE tissue sections of mouse liver or mouse kidney sectioned at various thicknesses (**Fig. 2A-M, Supplemental Fig**.**S5-11**).

**Figure 2:**
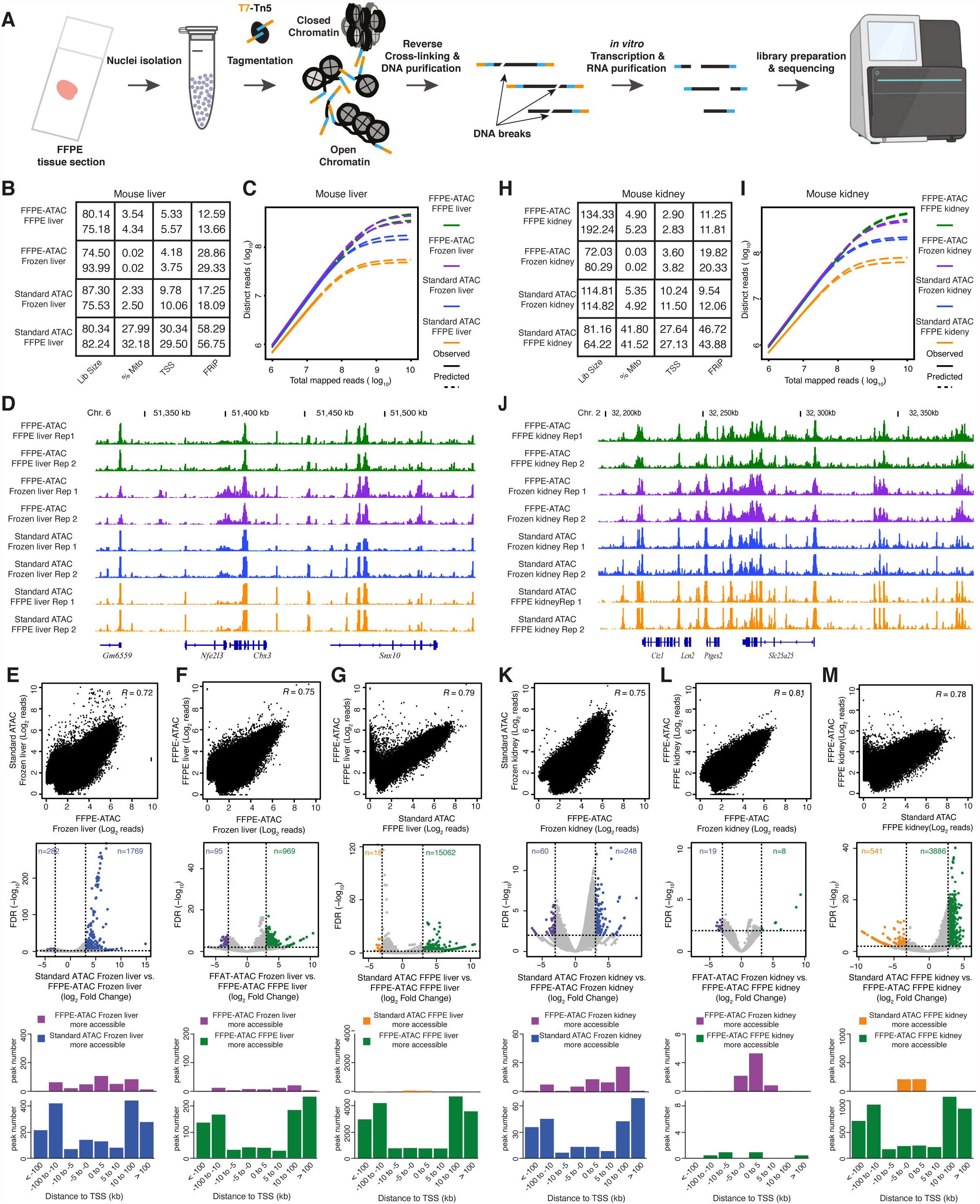
FFPE-ATAC decodes chromatin accessibility with low cell numbers obtained from FFPE tissue sections. (**A**), Workflow of FFPE-ATAC. (**B**), Quality control metrics of FFPE-ATAC on frozen mouse liver and FFPE mouse liver, and standard ATAC-seq on frozen mouse liver and FFPE mouse liver. Lib size = total sequencing reads of sequencing library (million); %Mito = percentage of mitochondria; TSS = enrichment score at transcription start sites (TSS); FRiP = fraction of reads in peaks. (**C, D**), Comparison of sequencing library complexity (**C**) and genome browser tracks (**D**) from FFPE-ATAC on frozen mouse liver and FFPE mouse liver, and standard ATAC-seq on frozen mouse liver and FFPE mouse liver. Chr. = Chromosome. (**E-G**), Comparison of chromatin accessibility from different conditions: standard ATAC-seq on frozen mouse liver vs. FFPE-ATAC on frozen mouse liver (**E**), FFPE-ATAC on frozen mouse liver vs. FFPE-ATAC on FFPE mouse liver (**F**), and FFPE-ATAC on FFPE mouse liver vs. standard ATAC-seq on FFPE mouse liver (**G**). Top: Genome-wide comparison of accessible chromatin regions. *R* = Pearson’s correlation. Middle: Differential peak analysis. FDR = false discovery rate. Bottom: Distribution of the more accessible regions from each condition across transcription start sites (TSS). (**H**), Quality control metrics of FFPE-ATAC on frozen mouse kidney and FFPE mouse kidney, and standard ATAC-seq on frozen mouse kidney and FFPE mouse kidney. Lib size = total sequencing reads of sequencing library (million); %Mito = percentage of mitochondria; TSS = enrichment score at transcription start sites (TSS); FRiP = fraction of reads in peaks. (**I, J**), Comparison of sequencing library complexity (**I**) and genome browser tracks (**J**) from FFPE-ATAC on frozen mouse kidney and FFPE mouse kidney, and standard ATAC-seq on frozen mouse kidney and FFPE mouse kidney. Chr. = Chromosome. (**K-M**), Comparison of chromatin accessibility from different conditions: standard ATAC-seq on frozen mouse kidney vs. FFPE-ATAC on frozen mouse kidney (**K**), FFPE-ATAC on frozen mouse kidney vs. FFPE-ATAC on FFPE mouse kidney (**L**), and FFPE-ATAC on FFPE mouse kidney vs. standard ATAC-seq on FFPE mouse kidney (**M**). Top: Genome-wide comparison of accessible chromatin regions. *R* = Pearson’s correlation. Middle: Differential peak analysis. FDR = false discovery rate. Bottom: Distribution of the more accessible regions from each condition across transcription start sites (TSS).

First, we cut a mouse liver into two parts, one part was frozen, and the other was prepared as an FFPE block (**Supplementary Fig. S5A**, see **Methods**). We performed FFPE-ATAC on nuclei purified from frozen mouse liver and FFPE mouse liver (**Supplemental Fig. S5A**, see **Methods**). Sequencing libraries obtained from frozen mouse liver by FFPE-ATAC had good genome-wide reproducibility (**Supplemental Fig. S5B**). The sequencing reads of the libraries were enriched at TSS (**Supplemental Fig. S5C**), but the TSS enrichment score was 1.5-2.5-fold lower than those of libraries obtained by standard ATAC-seq on frozen samples (**Fig. 2B**). However, the sequencing library complexity obtained from FFPE-ATAC on frozen mouse liver is much higher that obtained from standard ATAC-seq on frozen mouse liver (**Fig. 2C**). The reason for the lower complexity of standard ATAC-seq libraries compared with FFPE-ATAC libraries is that standard ATAC-seq is a PCR-based method, and it requires two correct pairs of Tn5 adaptor insertions (Buenrostro et al. 2013). One insertion event or unpaired Tn5 adaptor insertions from Tn5 tagmentation could not be amplified through PCR in standard ATAC-seq but could be captured with FFPE-ATAC. FFPE-ATAC on frozen mouse liver and standard ATAC-seq on frozen mouse liver exhibited high similarity at the level of chromatin accessibility at individual gene loci (**Fig. 2D)**, and in the distribution of sequence reads across the genome (**Supplemental Fig. S5D, S5E**). The two libraries also showed good genome-wide correlation (*R* = 0.72, **Fig. 2E)** and displayed a large number of overlapping ATAC peaks (53 043 overlapping peaks, **Supplemental Fig. S6A**). Some differential peaks are detected between FFPE-ATAC on frozen mouse liver and standard ATAC-seq on frozen mouse liver (Log_2_(fold change) > 3, p < 0.01) (Supplemental Code) (n = 262 in FFPE-ATAC, and n = 1789 in standard ATAC-seq, **Fig. 2E**), which indicates that there are potentially different technical biases between FFPE-ATAC and standard ATAC-seq. Our results suggested that FFPE-ATAC could accurately profile chromatin accessibility in frozen samples with higher library complexity than standard ATAC-seq. Next, compared the sequencing libraries obtained using FFPE-ATAC with FFPE mouse livers and frozen mouse livers. We found high similarity at the level of library complexity (**Fig. 2B**), TSS enrichment score (**Fig. 2C**), chromatin accessibility at individual gene loci (**Fig. 2D)**, and sequence read distribution across the genome (**Supplemental Fig. S5D, S5E**). There was also a good genome-wide correlation (*R* = 0.75, **Fig. 2F)**, and a large number of overlapping ATAC peaks (49530 overlapping peaks, **Supplemental Fig. S6B**). At the same time, we found that the TSS enrichment scores obtained by FFPE-ATAC on frozen mouse liver and FFPE mouse liver were similar to each other but 1.5- to 2.5-fold lower than the scores obtained by standard ATAC-seq on frozen mouse liver. This could be due to the different designs of FFPE-ATAC and standard ATAC-seq. Differential peak analysis showed that only 95 more accessible chromatin regions were captured from FFPE-ATAC on frozen mouse liver, but 969 more accessible chromatin regions were detected from FFPE-ATAC on FFPE mouse liver (**Fig. 2F, Supplemental Table S3**). The similar levels of library complexity obtained through FFPE-ATAC on FFPE mouse liver and FFPE-ATAC on frozen mouse liver and the very limited number (n = 95) of more accessible chromatin regions detected from FFPE-ATAC on frozen mouse liver suggest that FFPE-ATAC can potentially decode all accessible chromatin sites in the genome by rescuing broken DNA fragments in FFPE mouse liver. However, the FRiP from FFPE-ATAC on FFPE mouse liver (approximately 13%) was much lower than that from FFPE-ATAC on frozen mouse liver (approximately 29%) (**Fig. 2B**); this could be due to the harsh chemical treatments used during the preparation of FFPE samples. Finally, we compared the sequencing libraries obtained by FFPE-ATAC on FFPE mouse liver and by standard ATAC-seq on FFPE mouse liver, and found that the library complexity obtained from FFPE-ATAC was much higher than that obtained from standard ATAC-seq (**Fig. 2C**). Even though there is high similarity between FFPE-ATAC on FFPE mouse liver and standard ATAC-seq on FFPE mouse liver based on multiple comparisons (**Fig. 2C-D, 2G, Supplemental Fig. S6C**), we identified 15 062 more accessible chromatin regions in FFPE-ATAC on FFPE mouse liver, and these were mainly distributed in regions distal to TSS (**Fig. 2G**). However, only 18 more accessible chromatin regions were detected in standard ATAC-seq on FFPE mouse liver (**Fig. 2G**). We reasoned that if those large numbers of more accessible regions in FFPE-ATAC on FFPE mouse liver are located at sites of DNA breakage in FFPE mouse liver, no sequencing reads from those regions would be detected in libraries prepared from FFPE mouse liver by standard ATAC-seq. Indeed, among the more accessible regions identified through FFPE-ATAC on FFPE mouse liver, 71.83% (10819/15062) of those regions had no PCR amplicons in the libraries obtained by standard ATAC-seq on FFPE mouse liver (**Supplemental Fig. S6D, Supplemental Table S4**); this strongly indicates that FFPE-ATAC can be used to rescue accessible regions at DNA breakage sites in FFPE mouse liver samples. Taken together, our results demonstrate that the accessible chromatin profiles obtained using FFPE-ATAC on FFPE mouse liver are very similar to the accessible chromatin profiles in frozen mouse liver. The strategy used in FFPE-ATAC can thus rescue accessible regions that are lost due to DNA breaks when standard ATAC-seq of FFPE samples is used, resulting in greater library complexity and higher coverage of accessible chromatin profiles.

Second, following the same strategy that was used with mouse liver, we performed FFPE-ATAC on frozen mouse kidney and on FFPE mouse kidney, and conducted cross-comparisons among libraries prepared by FFPE-ATAC on FFPE mouse kidney, FFPE-ATAC on frozen mouse kidney, standard ATAC-seq on FFPE mouse kidney and standard ATAC-seq on frozen mouse kidney (**Fig. 2H-M, Supplemental Fig. S7, Supplemental Fig. S8, and Supplemental Table S5-8**). We also obtained high-quality FFPE-ATAC results from mouse FFPE kidneys (**Supplemental Fig. S7B-D**). The FFPE-ATAC on FFPE mouse kidney and that on frozen mouse kidney from the same mouse kidney also exhibited high similarity in library complexity (**Fig. 2I**), chromatin openness at the level of individual gene loci (**Fig. 2J**) and genomic features of ATAC peaks (**Supplemental Fig. S7D, S7E**). There was a good genome-wide correlation (*R* = 0.81, **Fig. 2 L**) and a large number of overlapping ATAC peaks (63 259 overlapping peaks, **Supplemental Fig. S8C**) in the results obtained from FFPE-ATAC on FFPE mouse kidney and FFPE-ATAC on frozen mouse kidney. A very limited number of differential peaks (n = 19 in FFPE-ATAC on frozen mouse kidney, n = 8 in FFPE-ATAC of FFPE mouse kidney, **Fig. 2 L**) between FFPE-ATAC on frozen mouse kidney and that on FFPE mouse kidney were identified, indicating that the chromatin profiles captured with FFPE-ATAC on FFPE mouse kidney are very similar to those captured with FFPE-ATAC on frozen mouse kidney. Differential peak analysis of FFPE-ATAC on FFPE mouse kidneys and standard ATAC-seq on FFPE mouse kidney showed that 3886 more accessible chromatin regions were decoded in FFPE-ATAC on FFPE mouse kidney (Supplemental Code), whereas only 541 more accessible chromatin regions were captured in standard ATAC-seq on FFPE mouse kidney (**Fig. 2 M**). For 61.65% (2396/3886) of the more accessible chromatin regions captured in FFPE-ATAC on FFPE mouse kidney, no sequencing reads were detected in those regions from libraries obtained by standard ATAC-seq on FFPE mouse kidney (**Supplemental Fig. S8D, Supplemental Table S8**). These results further demonstrate that FFPE-ATAC can profile accessible chromatin with better library complexity and rescue accessible regions at sites of DNA breakage in FFPE samples compared with standard ATAC-seq on FFPE samples. Analysis of FFPE-ATAC libraries generated from both FFPE mouse liver and FFPE mouse kidney revealed a large number of peaks that overlap with the peaks listed in the Encyclopedia of DNA Elements (ENCODE) mouse liver or kidney DNase-seq; there were 39 378 overlapping peaks for mouse liver (**Supplemental Fig. S9A**) and 64 612 overlapping peaks for mouse kidney (**Supplemental Fig. S9B**).

Third, we tested the sensitivity of FFPE-ATAC using various numbers of nuclei (ranging from 500 to 50 000) purified from FFPE mouse kidney tissue (**Supplemental Fig. S10A**, see **Methods**) Based on a comprehensive comparison of chromatin accessibility obtained using 50 000 nuclei, including TSS enrichment scores (**Supplemental Fig. S10B**), FRiP values (**Supplemental Fig. S10B**), library complexity (**Supplemental Fig. S10C**), genome-wide correlation (**Supplemental Fig. S10D-F**), and sequencing read distribution across the genome (**Supplemental Fig. S10G, S10H**), we concluded that FFPE-ATAC resulted in good accessible chromatin profiles of FFPE samples when as few as 500 nuclei were used.

Fourth, we determined the minimum thickness of FFPE tissue sections needed for the FFPE-ATAC by performing the FFPE-ATAC with 50 000 nuclei isolated from the 5-, 7-, and 10-µm thick mouse FFPE kidney tissue sections (**Supplemental Fig. S11A, S11B;** see **Methods**). The diameter of a mammalian cell nucleus is 6-10 µm (Webster et al. 2009), whereas the FFPE tissue sections used in routine clinical practice are 4-50 µm thick. We therefore investigated whether satisfactory FFPE-ATAC results could be obtained using FFPE tissue sections of different thicknesses. We found that the TSS enrichment score, library complexity and other parameters of the libraries obtained from mouse kidney FFPE-ATAC remained adequate when 5 µm thick tissue sections were used (**Supplemental Fig. S11C-F**, see **Methods**). However, FRiP values of FFPE-ATAC libraries obtained from 5-, 7-, and 10-µm-thick mouse FFPE kidney tissue sections, ranging from 2.4% to 7.5% (**Supplementary Fig. 11C**), were all lower than that of the FFPE-ATAC library obtained from a 20-µm-thick mouse FFPE kidney tissue section (∼11%). In addition, the total number of accessible peaks in the libraries prepared from these thin sections was much lower than the number of accessible peaks in the libraries prepared from 20-µm-thick mouse FFPE kidney tissue sections (**Supplemental Fig. S11G)**. Since the diameter of the mammalian nucleus is 6-10 µm (Webster et al. 2009), we reasoned that nuclei isolated from 5- to 10-µm-thick FFPE tissue sections contain a large proportion of nonintact nuclei. We suspected that the structure of the chromatin in nonintact nuclei could be affected during the isolation procedure, resulting in low-quality accessible chromatin profiles. Thus, we concluded that FFPE tissue sections with thickness greater than the diameter of nucleus should be used in FFPE-ATAC.

Taken together, accurate mapping of the accessible genome from mouse FFPE liver and kidney tissue sections demonstrates that FFPE-ATAC can be used to identify the accessible chromatin landscape using low number of cells obtained from FFPE tissue sections.

### Use of combination of FFPE-ATAC and H&E staining to decipher chromatin accessibility in a region of interest in FFPE tissue sections

Next, we deciphered chromatin accessibility in the mouse cerebellum by using hematoxylin and eosin (H&E) staining to identify mouse cerebellum in FFPE tissue sections of the mouse brain (**Fig. 3A-D**). H&E staining is a standard method that is used in clinical diagnostics to facilitate the assessment of tumor morphology and composition (Martina et al. 2011). We used H&E staining of a 5-µm-thick FFPE mouse brain tissue section to find the location of the cerebellum; we then isolated the cerebellar region from the immediately adjacent 20-µm-thick FFPE mouse brain tissue section, and purified the nuclei from the isolated cerebellum for use in FFPE-ATAC (**Fig. 3A, Supplemental Fig. S12A**, see **Methods**). The resulting FFPE-ATAC profiles of the mouse cerebellum had good TSS enrichment scores, FRiP values (**Supplemental Fig. S12B**), library complexity (**Supplemental Fig. S12C**) and genomic features (**Supplemental Fig. S12D, S12E**). The chromatin accessibility of regulatory elements of cerebellum-specific genes such as *Gabrb2*, was high (**Fig. 3B**). The technical replicates for FFPE-ATAC libraries from the cerebellum exhibited good reproducibility of genome-wide correlation, showing with numerous overlapping peaks (*R* = 0.86, 58 277 overlapping ATAC-seq peaks, **Fig. 3C**). Gene Ontology term enrichment and Kyoto Encyclopedia of Genes and Genomes (KEGG) pathway analysis for the top 10 000 FFPE-ATAC peaks identified major terms and pathways that clearly represent relevant gene pathways of the cerebellum(Sato et al. 2008) (**Fig. 3D, Supplemental Fig. S12F**).

**Figure 3:**
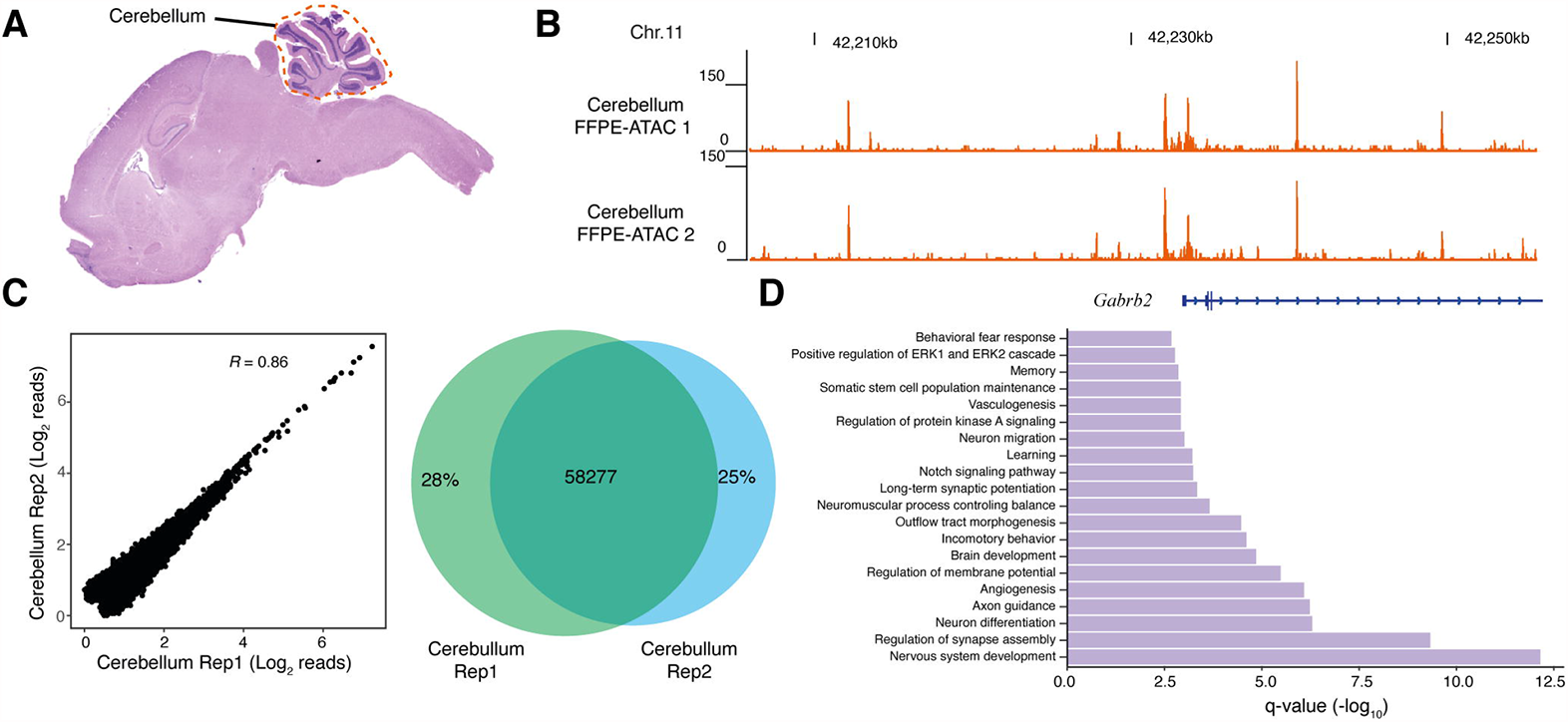
FFPE-ATAC decodes chromatin accessibility from the mouse cerebellum with the aid of H&E staining. (**A**), Hematoxylin and eosin staining (H&E) of a mouse FFPE brain tissue section, where the location of the cerebellum is illustrated with a dotted line. (**B**), Genome browser tracks of results from FFPE-ATAC analyses of isolated mouse FFPE cerebellum. Chr. = Chromosome. (**C**), Reproducibility of FFPE-ATAC analyses of mouse FFPE cerebellum. Left, the genome-wide correlation from the FFPE-ATAC reads. Right: the overlapping peaks from the FFPE-ATAC in the two technical replicates. *R* = Pearson’s correlation. (**D**), Enrichment of Gene Ontology terms for the top 10 000 FFPE-ATAC peaks for the mouse FFPE cerebellum.

### Application of FFPE-ATAC to clinically archived FFPE samples

Finally, we applied the FFPE-ATAC method to colorectal cancer (CRC) FFPE tissue sections from seven patients, including two cases of rectal cancer and five cases of colon cancer (**Fig. 4A-G, Supplemental Fig. S13A, S13B, Supplemental Table S9**). These CRC FFPE tissue blocks had been preserved for 6 to 10 years (**Supplemental Table S9**). The FFPE-ATAC libraries obtained from these CRC samples had good reproducibility (**Supplemental Fig. S13C, S13D**, *R* ranged from 0.86 to 0.97), good TSS enrichment scores (**Supplemental Fig. S14A**), library complexity (**Supplemental Fig. S14B**), and similar distributions of genomic features (**Supplemental Fig. S14C, S14D**). However, diverse ranges of FRiP, ranging from 5.78% to17.74%), was observed in the libraries from these clinical samples (**Supplemental Fig. S14A**); this could be due to variation in the procedures used for FFPE sample preparation. When we derived nonnegative matrix factorization (NMF) clusters using all the 7 CRC FFPE-ATAC peaks(Brunet et al. 2004), we found that two clusters were the best to characterize the 7 cases of CRC (**Fig. 4B, Supplemental Fig. S15A**); samples from three of the colon cancer patients were in cluster 1, while samples from the two rectal cancer patients and samples from the other two colon cancer patients were in cluster 2. The promoter regions of the CRC-specific gene marker *LRCH4* (Uhlen et al. 2015), were open in both cluster 1 and cluster 2 (**Fig. 4C**). Comparing the open chromatin sites within these two clusters, we identified 4 186 unique ATAC peaks for cluster 1 and 4 392 unique ATAC peaks for cluster 2 (fold change > 2, false discovery rate < 0.01, **Fig. 4D, 4E, Supplemental Fig. S15B, S15C, Supplemental Table S10, Supplemental Table S11, Supplemental Code**). We also found that the unique regulatory elements in these two clusters had similar genomic features (**Supplemental Fig. S15D, S15E**), but the ranking of transcription factors (TFs) enriched in the cluster-specific peaks are different between the two clusters (**Fig. 4F, 4G, Supplemental Table S12, Supplemental Table S13**); the top-ranking TFs for cluster 1 were ZIC1, TAL1, and NANOG, while the top-ranking TFs for cluster 2 were FOSL2, FOSL1 and JUN. It has been reported that AP-1 TFs play a dominant role in the progression of CRC(Ashida et al. 2005). We found 8 of the top 10 enriched TFs in cluster 2 are all from the AP-1 TF family (**Supplemental Fig. S15F**). A similarly high enrichment of AP-1 TF family members was not observed in cluster 1, likely reflecting a role of AP-1 TFs in some cases of CRC but not in others.

**Figure 4:**
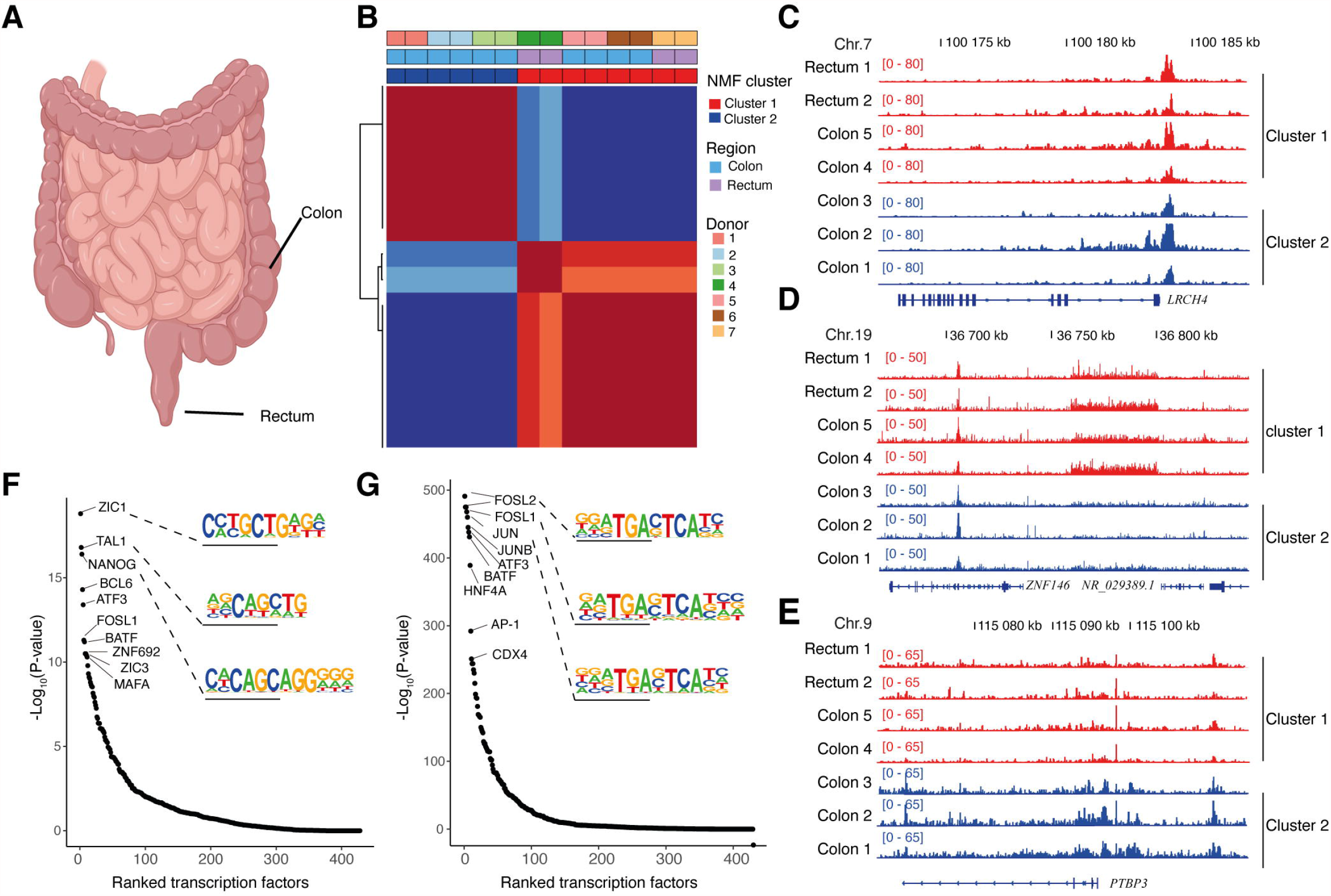
FFPE-ATAC decodes chromatin accessibility from clinical archived tumor samples. (**A**), Schematic image showing the location of human colorectal cancer (CRC) samples: colon and rectum. (**B**), Nonnegative matrix factorization (NMF) of chromatin accessibility with FFPE-ATAC from 2 cases of rectal cancer and 5 cases of colon cancer, identifying two clusters. (**C**), Regulatory elements of the CRC marker gene *LRCH4* are accessible in both clusters. (**D**), Representative gene loci that are more accessible in cluster 1, as seen from the differential FFPE-ATAC peaks. (**E**), Representative gene loci that are more accessible in cluster 2, as seen from the differential FFPE-ATAC peaks. (**F**), Ranked transcription factors significantly enriched in the specific regulatory elements from cluster 1. (**G**), Ranked transcription factors significantly enriched in the specific regulatory elements from cluster 2.

In summary, FFPE-ATAC allows the profiling of chromatin accessibility in specific regions of interest when combined with the use of H&E staining to identify the cell analyzed. This approach serves to identify unique distal regulatory elements and TF enrichments using low numbers of nuclei prepared from single clinically archived FFPE tissue sections preserved for extended periods of time.

## DISCUSSION

FFPE tissue samples represent a large source of materials for epigenetic analysis in both basic research and clinical translational studies(Gaffney et al. 2018), but such samples have not been widely used in chromatin studies to date due to the lack of sufficiently sensitive techniques. The broad application of ATAC-seq in biomedical research has offered us a potential strategy for profiling chromatin accessibility in FFPE samples with high sensitivity (Buenrostro et al. 2013; Buenrostro et al. 2015; Cusanovich et al. 2015; Chen et al. 2016; Corces et al. 2017; Corces et al. 2018). However, the presence of DNA damage in FFPE samples hampers the direct application of standard ATAC-seq to these samples(Chin et al. 2020). Using an optimized nuclei isolation protocol with FFPE tissue sections, we showed that transposase-mediated technology, ATAC-seq, could be applied to FFPE samples. However, standard ATAC-seq libraries of FFPE samples have lower library complexity and a smaller proportion of long DNA fragments, and lack a proportion of accessible chromatin sites compared with libraries obtained by standard ATAC-seq on frozen samples. To increase library complexity and rescue accessible regions that are lost in standard ATAC-seq on FFPE samples, we developed FFPE-ATAC, which used a combination of Tn5-mediated transposition and T7 IVT to decode chromatin accessibility in FFPE tissues. We demonstrated that the accessible chromatin profiles derived from FFPE samples by FFPE-ATAC are very similar to the accessible chromatin profiles of frozen samples. We learned that the TSS enrichment scores obtained after FFPE-ATAC of frozen samples and FFPE samples are similar to each other but 1.5-2.5-fold lower than those obtained through standard ATAC-seq of frozen samples; this could be due to the different designs of FFPE-ATAC and standard ATAC-seq. At the same time, we observed that the proportion of sequencing signals in peaks from FFPE-ATAC of FFPE samples was lower than the proportion in peaks from FFPE-ATAC of frozen samples but fell in a range similar to that in peaks from standard ATAC-seq of frozen samples; this could be due to the use of harsh chemical treatments during preparation of the FFPE samples. FFPE-ATAC is more labor-intensive than the more simply designed standard ATAC-seq method. However, through use of the FFPE-ATAC strategy, it was possible to rescue many of the accessible regions that are lost in standard ATAC-seq on FFPE samples, resulting in better library complexity. The better library complexity and higher coverage of accessible chromatin profiles that can be obtained using FFPE-ATAC on FFPE samples compared with standard ATAC-seq of FFPE samples will be valuable for accessible chromatin profiling of clinically archived FFPE materials.

We demonstrate here that FFPE-ATAC is a robust tool that can be used to decode chromatin accessibility with high sensitivity using 500-50 000 nuclei prepared from single FFPE tissue sections. The use of a combination of FFPE-ATAC and H&E staining to decipher chromatin accessibility in a region of interest in FFPE tissue sections and successful profiling of disease-associated chromatin regulation from the clinically archived human colorectal cancer FFPE samples make FFPE-ATAC a powerful tool for use in preclinical studies and precision medicine. In addition, FFPE-ATAC can potentially be used to extend our current understanding of the cancer epigenome and in pathological diagnosis through combination of other omics data obtained from the same FFPE materials with clinicopathological records. FFPE-ATAC can find broad applications both in basic research and in clinical settings. In the future, it will be of great interest to extend the resolution of FFPE-ATAC to the single-cell level.

## METHODS

### Nuclei isolation from FFPE tissue sections

Mouse FFPE kidney, liver, and brain tissue blocks were sectioned into 5-, 7-, 10-, and 20-*µ*m thick section using a microtome. The human CRC samples were cut into 10 *µ*m thick section. One curved tissue section was deparaffined with 1ml of xylene (HistoLab, 02070) 5mins, twice. Rehydration was done by sequential ethanol washes, started with 100% ethanol 5 min twice, 95%, 70%, 50%, 30% ethanol, 5 min each. After deparaffinization and rehydration, tissue was washed with 1 ml water, then 1ml 0.5mM CaCl_2_ (Alfa Aesar, J63122). Then the tissue was subjected to microdissection under a stereo microscope first, then centrifuged at 3000g for 10 mins. After centrifugation, supernatant was removed and 1ml enzymatic cocktail (3mg/ml of Collagenase (Sigma-Aldrich, C9263) and 300 U/ml of hyaluronidase (Merk Millipore, HX0154-1)) was added to the tissue pellet. Then the mixture was incubated at 37°C for 16 hours by adding 100*µ*g of Ampicillin (Serva, 69-52-3) and 50*µ*g of sodium azide (Merck Millipore,26628-22-8). After the enzyme digestion, 400*µ*l NST buffer (146mM NaCl (Invitrogen, 00648496), 10mM Tris pH 7.8 (Invitrogen, 15568-025), 1mM CaCl_2_ (Alfa Aesar, J63122), 21mM of MgCl_2_ (Invitrogen, AM9530G), 0.05% BSA (Miltenyi Biotech MACS, 130-091-376), 0.2% Igepal CA-360 (Sigma-Aldrich, 13021-50)) was added to the mixture, and the tube was centrifuged at 2800g for 10 minutes. After the centrifugation, the supernatant was aspirated and discarded, then the pellet was resuspended in 800 *µ*l NST buffer containing 0.1% DNase free RNase A (Thermo Scientific, EN0531), and 10% fetal bovine serum (Life Technologies,10108-105). The mixture was passed through the 27G needle syringe 30 times and filtered with a 30 *µ*M filter (MiltenyiBiotechMacs,130-098-458). Then the passed-through nuclear suspension was centrifugation at 2800 g for 10 minutes, and the nuclei were resuspended in PBS, checked and counted.

### Nuclei isolation from the Mouse cerebellum FFPE tissue section

The mouse cerebellum area was identified with H&E staining from the adjacent tissue sections, and labeled with marker pen under the stereo microscope. The tissues from the cerebellum area were moved to the Eppendorf tube and the nuclei isolation from the selected area was performed with the protocol stated as above.

### Human CRC sample collections and FFPE block preparation

The regional ethical research committee at the Uppsala University approved the study (Dnr 2015/419 and 2018/490). The FFPE tissue blocks of CRC were prepared at the Dept. of Clinical Pathology, Uppsala University Hospital, Uppsala, Sweden, according to standard procedures. Briefly, tissue from surgical specimens of colon and rectal samples were fixed in buffered formalin for 24-72 hours. The pieces were then examined by a pathologist, excised and placed in plastic cassettes. The fixed tissue was then dehydrated in an automated system (Tissue-Tek® VIP®) where the tissue was immersed in ethanol of varying concentrations (70%, 95%, 99.5%) followed by xylene and finally paraffin (Histowax™, Histolab) over a period of approximately 12 hours. Finally, the paraffin embedded tissue piece was oriented in a cassette, liquid paraffin was poured over it and allowed to set, forming the FFPE block. The FFPE block was then sectioned on a microtome at a thickness of 10-*µ*m.

### FFPE-ATAC on FFPE tissue and frozen tissue

For FFPE-ATAC on FFPE tissue, 500-50 000 isolated FFPE nuclei were used in each FFPE-ATAC reaction, where nuclei were isolated following nuclei isolation protocol stated in section of nuclei isolation from FFPE tissue sections. For FFPE-ATAC on frozen tissue, 50 000 isolated nuclei were used in each reaction, where nuclei were isolated following nuclei isolation protocol in section of standard ATAC-seq on frozen tissue. In brief, nuclei were counted using the cell counter and pelleted at 2800g for 10 minutes at room temperature. 50*µ*l of lysis buffer (10mM Tris-HCl pH7.4 (Invitrogen, 15567-027), 10mM NaCl (Invitrogen, AM9759), 3mM MgCl_2_ (Invitrogen, AM9530G), 0.1% Igepal CA-360 (Sigma-Aldrich, 13021-50)) was added to the nuclei pellet and the nuclei suspension was immediately centrifuged at 2800g for 10 minutes at room temperature. After the supernatant was discarded, the nuclei pellet was resuspended in 50 *µ*l of transposase master mixture (25*µ*l of 2X TD buffer (20mM Tris-HCl pH 7.6 (Invitrogen, 15568-025), 10mM MgCl_2_ (Invitrogen, AM9530G), 20%Dimethyl Formamide), 22.5 *µ*l Nuclease free water (Invitrogen, AM9932) and 2.5*µ*l of 2*µ*M T7-Tn5), then incubated at 37°C for 30 minutes. After the incubation, 50 *µ*l of 2X reverse crosslinking solution (100mM Tris-Cl PH 8.0 (Invitrogen, 15568-025), 2mM EDTA (Invitrogen, AM9290G), 2% SDS (Invitrogen, 15553-035), 0.4M NaCl (Invitrogen, AM9759)) and 10ng/ul Proteinase K (Thermo Scientific, EO0491) was directly added into the tagmentation reaction mixture, then mixture was incubated overnight at 65°C with 1200rpm shaking. Next day, the incubation mixture is purified with MinElute PCR Purification kit (QIAGEN, 28004) and DNA is eluted in 20*µ*l of elution buffer, then 20 *µ*l of 2X PCR master mix (New England Biolabs, New England Biolabs, M0541S) was added to the samples. The mixture was incubated in a thermocycler at 72°C for 5 minutes. The sample was first purified with MinElute PCR Purification kit (QIAGEN, 28004), then re-purified with SPRI beads with 1:1 ratio (Beckman Coulter, B23317), and eluted in 25 *µ*l of water (Invitrogen, AM9932).

Next, *In Vitro* Transcription (IVT) was performed with T7 high yield RNA synthesis kit (New England Biolabs, E2040S). The RNA from the IVT was purified using TRIzol first (Ambion, 15596026), then ZYMO RNA Clean & Concentration kit (Zymo, R1013). Next, 1 *µ*l DNase I (New England Biolabs, M0303L) was added into the RNA and the mixture was incubated for 15 minutes at 37°C. The RNA was purified with the ZYMO RNA Clean & Concentration kit (Zymo, R1013) again and eluted in 15 *µ*l of nuclease-free water. The IVT RNA is transferred into cDNA with random primers with SMART MMLV kit by following the manufactory protocol (TaKaRa, 639524). 100ng RNA was used for each library preparation. In brief, the mixture was incubated at 42°C for 60 minutes and 70°C for 15 minutes, then 2 *µ*l of RNase H buffer and 0.2 *µ*l RNase H enzyme (Thermo Scientific, EN0201) was added and incubated at 37°C for 20 minutes. The cDNA was purified using RNA XP beads purification with 1:1.8x ratio of sample to beads (Beckman Coulter, A63987) and eluted in 24.5 *µ*l water. Next, the cDNA was converted to double-stranded DNA with pre-PCR (98°c for 10 seconds, 63°C for 30 seconds, 72°C for 1 minute, 10°C hold in one cycle) by adding 25 *µ*l of 2X PCR master mix (New England Biolabs, M0541S) and 0.8 *µ*l of Ad 2.X reverse primer. Then sample was purified using MinElute PCR Purification kit (QIAGEN, 28004) and eluted in 20 *µ*l water.

The sequencing library was prepared with standard Tn5 tagmentation. In short, the double-strand DNA samples were subjected to the tagmentation by adding 25 *µ*l of 2X TD-Buffer (20mM Tris-HCl pH 7.6 (Invitrogen, 15567-027), 10mM MgCl_2_ (Invitrogen, AM9530G), 20% Dimethyl Formamide), 0.5 *µ*l 2uM standard Tn5, 4.5 *µ*l nuclease-free water (Invitrogen, AM9932) and incubated at 55°C for seven minutes, then samples were purified using Qiagen MinElute PCR Purification kit (QIAGEN, 28004) and eluted in 20 *µ*l elution buffer. The library amplification PCR was performed by adding 25 ul of 2X PCR master mix, 0.4 *µ*l of barcodes forward primer i5 25 *µ*M, 0.4 *µ*l of barcodes reverse Primer i7 25 *µ*M, 4.2*µ*l of nuclease-free water to the sample, with the following PCR protocol (72 °C 5mins first, 20 cycles of 98°C for 10 seconds, 63°C for 30 seconds, 72°C for 1 minute), then sample was purified using Qiagen MinElute PCR Purification kit (QIAGEN, 28004) and eluted in 20 *µ*l. At last, the DNA library with the length of 220-1000bp was selected with PAGE gel purification for sequencing. The FFPE-ATAC libraries were sequenced on Illumina NovaSeq 6000, and at least 40 million 150bp single end or paired end sequencing reads were generated for each library.

### Hyperactive Tn5 transposase production

Hyperactive Tn5 was produced as previously described(Picelli et al. 2014). In brief, pTXB1-Tn5 plasmid (Addgene, 60240) was introduced into T7 Express LysY/Iq *E. coli* strain (NEB, C3013). 10 ml of overnight cultured *E. coli* was inoculated to 500 ml LB medium. After incubation for 1.5 hrs at 37°C, bacteria was incubated about 2.5 hrs at room temperature. When the OD600 = 0.9, Tn5 protein was induced by adding 0.25 mM IPTG for 4 hrs. *E. coli* pellet was resuspended in lysis buffer (20 mM HEPES-KOH pH 7.2, 0.8 M NaCl, 1mM EDTA, 10% glycerol, 0.2% Triton X-100, complete proteinase inhibitor (Roche,11697498001)) and lysed by sonication. 10% PEI was added to supernatant of lysate to remove bacterial genomic DNA. 10 ml chitin resin (NEB, S6651L) was added to the supernatant and incubated with rotating for 1 hr at 4°C. The resin washed by lysis buffer extensively. In order to cleave Tn5 protein from intein, lysis buffer containing 100 mM DTT was added to the resin and stored in 4°C. After 48 hrs, protein was eluted by gravity flow and collected in 1ml fractions. 1 *µ*l of each fraction was added to detergent compatible Bradford assay (Thermo Fisher Scientific, 23246) and peaked fractions were pooled and dialyzed against 2X dialysis buffer (100 mM HEPE-KOH pH7.2, 0.2 M NaCl, 0.2 mM EDTA, 2 mM DTT, 0.2% Triton X-100, 20% glycerol). Dialyzed Tn5 protein was concentrated by using ultracel 30-K column (Millipore, UFC903024) and the quantity of Tn5 was measured by Bradford assay and visualized on NuPAGE Novex 4-12% Bis-Tris gel (Thermo Fisher Scientific, NP0321) followed by Coomassie blue staining.

### T7-Tn5 and Tn5 adaptor sequences

The oligonucleotides for Tn5 and T7-Tn5 transposase adaptor were synthesized at INTERGATED DNA TECHNOLOGIES (IDT), and the sequences of oligonucleotide are as follows:

Tn5MErev, 5’-[phos]CTGTCTCTTATACACATCT-3’;

T7-Tn5ME: 5′/CAT GAG ATT AAT ACG ACT CAC TAT AGG GAG AAG ATG TGT ATA AGA GAC AG-3′;

Tn5ME-A: 5′ - TCGTCGGCAGCGTCAGATGTGTATAAGAGACAG-3′;

Tn5ME-B: 5′-GTCTCGTGGGCTCGGAGATGTGTATAAGAGACAG-3′.

### PCR primer sequences

The PCR primers were synthesized at INTERGATED DNA TECHNOLOGIES (IDT), and the sequences of primers were used by referring to the previous report (Buenrostro et al. 2015).

### Tn5 and T7-Tn5 transposase assembly

The assembly of Tn5 and T7-Tn5 transpossase were performed as described (Picelli et al. 2014). Briefly, oligonucleotides (T7-Tn5ME, Tn5MErev, Tn5ME-A, Tn5ME-B) were resuspended in water to a final concentration of 100 *µ*M each. Equimolar amounts of Tn5MErev/Tn5ME-T7, Tn5MErev/Tn5ME-A and Tn5MErev/Tn5ME-B were mixed in separate 200 *µ*l PCR tubes. These oligos mixtures were denatured on a thermocycler for 5 min at 95°C and cooled down slowly on the thermocycler by turning off the thermocycler. The T7-Tn5 transposase was assembled with the following components: 0.25 vol Tn5MErev/Tn5ME-T7 (final concentration of each double-strand oligo is now 50 µM each), 0.4 vol glycerol (100% solution), 0.12 vol 2X dialysis buffer (100 mM HEPES-KOH at pH 7.2, 0.2 M NaCl (Invitrogen, AM9759), 0.2 mM EDTA (Invitrogen, AM9290G), 2 mM DTT, 0.2% Triton X-100 (Sigma-Aldrich, T8787), 20% glycerol (Sigma-Aldrich, G9012-500)), 0.1 vol SL-Tn5 (50 *µ*M), 0.13 vol water. The reagents were mixed thoroughly but gently, and the solution was left on the bench at room temperature for 1 h to allow annealing of oligos to Tn5. The Tn5 transposase was assembly with same procedure as T7-Tn5 transposase but with following oligos: 0.25 vol Tn5MErev/Tn5ME-A and 0.25 vol Tn5MErev/Tn5ME-B.

### T7-Tn5 transposase activity assay

The activity of the assembled T7-Tn5 and Tn5 transposase was checked as described below. The mixture of 10*µ*l of 2X TD buffer (20mM Tris-HCl pH 7.6 (Invitrogen, 15568-025), 10mM MgCl_2_ (Invitrogen, AM9530G), 20%Dimethyl Formamide), 50ng human genomic DNA (Promega, G304A), 2*µ*M assembled T7-Tn5 transposase or Tn5 transposase was incubated at 55 °C for 7minutes. After incubation, the mixture was purified by Qiagen MinElute PCR Purification kit (QIAGEN, 28004) and eluted in 10*µ*l of elution buffer. Then eluted DNA was mixed with 2 *µ*l 6X loading dye and ran on a 1.2% agarose gel to check the length distribution of the DNA.

### Standard ATAC-seq on FFPE samples

50, 000 isolated FFPE nuclei (mouse liver and moue kideny) were used in each reaction following standard ATAC-seq protocol as previous reported(Buenrostro et al. 2013). The reverse-crosslinking was used after Tn5 tagmentation following the protocol of ATAC-seq in fixed cells (Chen et al. 2016). Briefly, 50,000 cells were centrifuged 500 g 5 min at room temperature. The cell pellet was resuspended in 50 µl lysis buffer (10 mM Tris-Cl, pH 7.4, 10 mM NaCl, 3 mM MgCl2, 0.01% Igepal CA-630) and centrifuged immediately 500 g for 10 min at 4 °C. The cell pellet was resuspended in 50 µl transposase mixture (25 µl 2X TD buffer, 22.5 µl dH2O and 2.5 µl Tn5 transposase) and incubated at 37 °C 30 min. After the transposase reaction, a reverse crosslink solution was added (with final concentration of 50 mM Tris-Cl, 1mM EDTA, 1% SDS, 0.2M NaCl, 5 ng/ml Proteinase K) up to 200 µl. The mixture was incubated at 65 °C with 1000 rpm shaking in a heat block overnight, then purified with Qiagen Mini-purification kit and eluted in 10 µl Quiagen EB elution buffer. Sequencing libraries were prepared following the original ATAC-seq protocol (Buenrostro et al. 2013).

### Standard ATAC-seq on frozen samples

Single nuclei were isolated from frozen tissue with Dounce homogenization by following the nuclei isolation protocol in Omni-ATAC(Corces et al. 2017). In brief, green bean size frozen tissue incubated in the ice-cold 800 *µ*l of 1X homogenization unstable buffer(5 mM CaCl2 (Alfa Aesar, J63122), 3 mM Mg(Ac)2 (Sigma-Aldrich, M5661), 10 mM Tris pH 7.8 (Invitrogen, 15568-025), 0.01667 mM PMSF (Sigma-Aldrich, P7626), 0.1667 mM β-mercaptoethanol (Sigma-Aldrich, M-6250), 320 mM Sucrose (Sigma-Aldrich, 84097-250), 0.1mM EDTA (Invitrogen, AM9290G), 0.1% Igepal CA-630 (Sigma-Aldrich, 13021-50)) for 5 minutes on ice.. Tissue was homogenized through 10 strokes with a loose pestle and 20 strokes with a tight pestle, then 400 *µ*l of the homogenized sample was mixed with 400 *µ*l of 50% OptiPrep Density Gradient Medium (Sigma-Aldrich, D1556-250), to make a final concentration of 25% of OptiPrep Density Gradient Medium (Sigma-Aldrich, D1556-250) with homogenized tissue. After preparation of tissue mixture, a fresh 2ml low binding vial was taken and layered 35% of OptiPrep Density Gradient Medium (Sigma-Aldrich, D1556-250), 29% of OptiPrep Density Gradient Medium (Sigma-Aldrich, D1556-250), and 25% of OptiPrep Density Gradient Medium (Sigma-Aldrich, D1556-250) mixed with the sample, on the top of each other. The layered vial was centrifuged at 3000g for 20 minutes at 4°C. After gradient centrifugation, the top 1300 *µ*l was discarded and the 200 *µ*l of the nuclei region was carefully collected in a fresh vial. Then 800 *µ*l of ice cold PBS was added and centrifuged at 500g for 10 mins, followed by resuspended in ice cold 500 *µ*l PBS. 50,000 nuclei were used for each reaction and prepared library by using standard ATAC protocol as stated in the section of standard ATAC-seq on FFPE samples (Buenrostro et al. 2013). The components for the solutions are as follows: 6X Homogenization Buffer Stable Master Mix: 30mM CaCl_2_ (Alfa Aesar, J63122), 18mM Mg(Ac)2, 60mM Tris-HCl pH 7.8 (Invitrogen, 15568-025); 6X Homogenization Buffer Unstable Solution: 6XHomogenization Buffer Stable Master Mix, 0.1mM PMSF, 1mM β-mercaptoethanol (Sigma-Aldrich, M-6250); 1X Homogenization Buffer Unstable Solution: 1XHomogenization Buffer Stable Master Mix, 320mM Sucrose (Sigma-Aldrich, 84097-250), 0.1mM EDTA (Invitrogen, AM9290G), 0.1% Igepal CA-360 (Sigma-Aldrich, 13021-50); 50% OptiPrep Density Gradient Medium (Sigma-Aldrich, D1556-250) Solution: 1XHomogenization Buffer Stable Master Mix, 50% OptiPrep Density Gradient Medium (Sigma-Aldrich, D1556-250)) Solution; 29% OptiPrep Density Gradient Medium (Sigma-Aldrich, D1556-250) Solution: 1XHomogenization Buffer Stable Master Mix, 160mM sucrose, 29% OptiPrep Density Gradient Medium (Sigma-Aldrich, D1556-250) Solution; 35% OptiPrep Density Gradient Medium (Sigma-Aldrich, D1556-250) Solution: 1XHomogenization Buffer Stable Master Mix, 160mM sucrose (Sigma-Aldrich, 84097-250), 35% OptiPrep Density Gradient Medium (Sigma-Aldrich, D1556-250) Solution. The ATAC-seq libraries were sequenced on Illumina NovaSeq 6000, and at least 20 million 150bp paired-end sequencing reads were generated for each library.

### Genomic DNA purification from frozen and FFPE tissue nuclei

For FFPE-ATAC samples, single nuclei were isolated following nuclei isolation protocol stated in section of nuclei isolation from FFPE tissue sections. For frozen samples, nuclei were isolated following nuclei isolation protocol in section of standard ATAC-seq on frozen tissue. For genomic DNA purification, 1 million isolated nuclei were spined down at 3000 g for 10 mins, then resuspended with 100 *µ*l of lysis buffer (50 mM Tris-HCl pH=7.5 (Invitrogen, 15567027), 1 mM EDTA (Invitrogen, AM9260G), 1% SDS (Invitrogen, 1553-035), 200 mM NaCl (Invitrogen, AM9759) and 200 *µ*g/mL Proteinase K (Thermo Scientific, EO0491). Nuclei suspension was incubated at 65 °C with 1200 rpm shaking in a heat block overnight. On the next day, the mixture was purified with Qiagen MiniElute Purification kit (QIAGEN, 28004) and eluted in 20 *µ*l of elution buffer. Purified genomic DNA was measured, and run on a 1.5% agarose gel (Lonza, 50004) to check size distribution.

### Animals

The mouse brain, liver, and kidney tissues were from the 8-week-old Mice FVBN mice, housed in individually ventilated cages (3-5 animals per cage) in accordance with Uppsala University regulations on mice with appropriate organic bedding, paper house enrichments, food and water *ad libitum* and 12/12-hour light/dark cycle. All experiments were performed in accordance with national guidelines and regulations, and with the approval of the animal care and use committees at Uppsala University.

### Mouse Tissue collection

8-week-old Mice were sacrificed via inhalation euthanasia, and mouse organs (brains, livers and kidneys) were collected. For frozen sample, livers and kidneys were snap-frozen on dry ice and stored at –80 °C. For FFPE sample, mouse brains, livers and kidneys were fixed with formalin overnight, and then washed with phosphate-buffered saline (PBS) and kept in 70% ethanol for paraffin embedding. Fixed mouse brains, livers and kidneys were routinely processed, and paraffin embedded.

### Primary data processing for the FFPE-ATAC and standard ATAC-seq

All scripts (available in Supplemental Code) are deposited in the following link: https://github.com/pengweixing/FFPE-ATAC.For sequencing libraries of FFPE-ATAC, the T7 promoter sequences and Tn5 transposase sequences from the Illumina single end sequencing reads were trimmed using cutadapt software with slightly modifications (Martin 2011) and in-house script, which was deposited in following link: https://github.com/pengweixing/FFPE-ATAC. For sequencing libraries of standard ATAC-seq, the Tn5 transposase sequences from the Illumina paired end sequencing reads were trimmed with in-house script. After the adaptor trimming, the sequencing reads were mapped to the reference genome (mm9 or hg19) with Bowtie 2 using parameters -very sensitive(Langmead et al. 2009). The duplicate reads were removed with Picard v1.79 (http://picard.sourceforge.net). The mapping for FFPE-ATAC on FFPE samples, FFPE-ATAC on frozen samples, standard ATAC-seq on FFPE samples and standard ATAC-seq on frozen samples was all performed with same parameters, thus, using GRCh38 and GRCm38 (mm10) as refence genome for mapping would not significantly affect the conclusions. SAMtools v1.9 software was used to sort and filter BAM files(Li et al. 2009). The bigWig file was generated from BAM file using deepTools v3.5 software with the option “bamCoverage”(Ramirez et al. 2014). The TSS enrichment score was calculated using deepTools with the option “computeMatrix”(Ramirez et al. 2014). The peak calling was performed using MACS2 in the parameters of -q 0.01 -nomodel -shift 0 (Zhang et al. 2008). The read counts within peaks for each sample were calculated using BEDTools v2.29.2 with the option “multicov”(Quinlan and Hall 2010). Genomic annotation and distance of peaks relative to TSS were calculated using ChIPseeker R package (Yu et al. 2015). Sequencing library complexity was calculated using Preseq v3.1.2 (Daley and Smith 2014). Differential peak analysis was performed with DESeq2 software (Love et al. 2014) and differential peaks were filtered with Log_2_ (fold change) >3 and false discovery rate <0.01. The insert size distribution for nucleosome-free region and mononucleosome were calculated using ATACseqQC package (Ou et al. 2018). The sequencing coverage was visualized in the Integrative Genomics Viewer (IGV) (IGV) (Thorvaldsdottir et al. 2013). Transcriptional factors enrichments were performed using HOMER v4.11 with “findMotifsGenome” tool (Heinz et al. 2010). The gene annotation was analyzed using ChIPseeker package(Yu et al. 2015). The Gene Ontology (GO) and Kyoto Encyclopedia of Genes and Genomes (KEGG) analysis were performed with DAVID(Huang et al. 2007).

### Differential peak analysis of CRC FFPE-ATAC

The Nonnegative matrix factorization (NMF) method(Brunet et al. 2004) was used to cluster the 7 cases of CRC FFPE-ATAC with default algorithm. The differentially FFPE-ATAC peaks from two clusters of CRC were identified with DEseq2(Love et al. 2014), following the parameter of fold-change > 2 and false discovery rate < 0.01. HOMER was used to calculate the significant transcriptional factors enrichment from the differentially FFPE-ATAC peaks(Heinz et al. 2010).

### ENCODE DNase-seq data

8-week-old mouse liver and kidney ENCODE DNase-seq data were downloaded from NCBI GEO with accession numbers: GSM1014195 (liver) and GSM1014193 (kidney).

## DATA ACCESS

All raw and processed sequencing data generated in this study have been submitted to the NCBI Gene Expression Omnibus (GEO; https://www.ncbi.nlm.nih.gov/geo/) under accession number GSE163306.

## ACKNOWLEDGMENTS

We thanks for the critical comments of our manuscript from Dr. Ulf Landgren. This work is supported by grants to X.C. from the Swedish Research Council (VR-2016-06794, VR-2017-02074.), Åke Wibergs stiftelse (M20-0007), Beijer Foundation, Jeassons Foundation, Petrus och Augusta Hedlunds Stiftelse, Göran Gustafsson’s prize for younger researchers, Vleugel Foundation, and Uppsala University). Part of this work was facilitated by the Protein Science Facility at Karolinska Institute, Stockholm.

## AUTHOR CONTRIBUTIONS

F. J. W., T. S. and X. C. conceived and designed the study. H. Z., V. K. P., M. Z., L. M., L. Z., G. R. performed experiments. P.X., and H. Z. performed all the data mining in the study. X. C. wrote the manuscript with input from all authors. X. C. supervised all aspects of this work.

