## Supplementary_FigS1_S15 for "Profiling chromatin accessibility in formalin-fixed paraffin-embedded samples"

^3^ Contributed equally

**Table of contents**

Pages 2-21: Supplemental Figures S1-S15 & legends

**Additional File**:

Supplemental Table S1: Lst of differential accessible chromatin peaks between standard ATAC-seq on FFPE mouse liver and standard ATAC-seq on frozen mouse liver.

Supplemental Table S2: List of differential accessible chromatin peaks between standard ATAC-seq on FFPE mouse kidney and standard ATAC-seq on frozen mouse kidney.

Supplemental Table S3: List of differential accessible chromatin peaks between FFPE-ATAC on frozen mouse liver and standard ATAC-seq on frozen mouse liver.

Supplemental Table S4: List of differential accessible chromatin peaks between FFPE-ATAC on frozen mouse liver and FFPE-ATAC on FFPE mouse liver.

Supplemental Table S5: List of differential accessible chromatin peaks between standard ATAC-seq on FFPE mouse liver and FFPE-ATAC on FFPE mouse liver.

Supplemental Table S6: List of differential accessible chromatin peaks between FFPE-ATAC on frozen mouse kidney and standard ATAC-seq on frozen mouse kidney.

Supplemental Table S7: List of differential accessible chromatin peaks between FFPE-ATAC on frozen mouse kidney and FFPE-ATAC on FFPE mouse kidney.

Supplemental Table S8: List of differential accessible chromatin peaks between standard ATAC-seq on FFPE mouse kidney and FFPE-ATAC on FFPE mouse kidney.

Supplemental Table S9: Sample Information of human colorectal cancer (CRC) FFPE tissue.

Supplemental Table S10: Differential peaks for cluster 1 from FFPE-ATAC of CRC.

Supplemental Table S11: Differential peaks for cluster 2 from FFPE-ATAC of CRC.

Supplemental Table S12: List of significant enriched TFs in the cluster 1.

Supplemental Table S13: List of significant enriched TFs in the cluster 2.

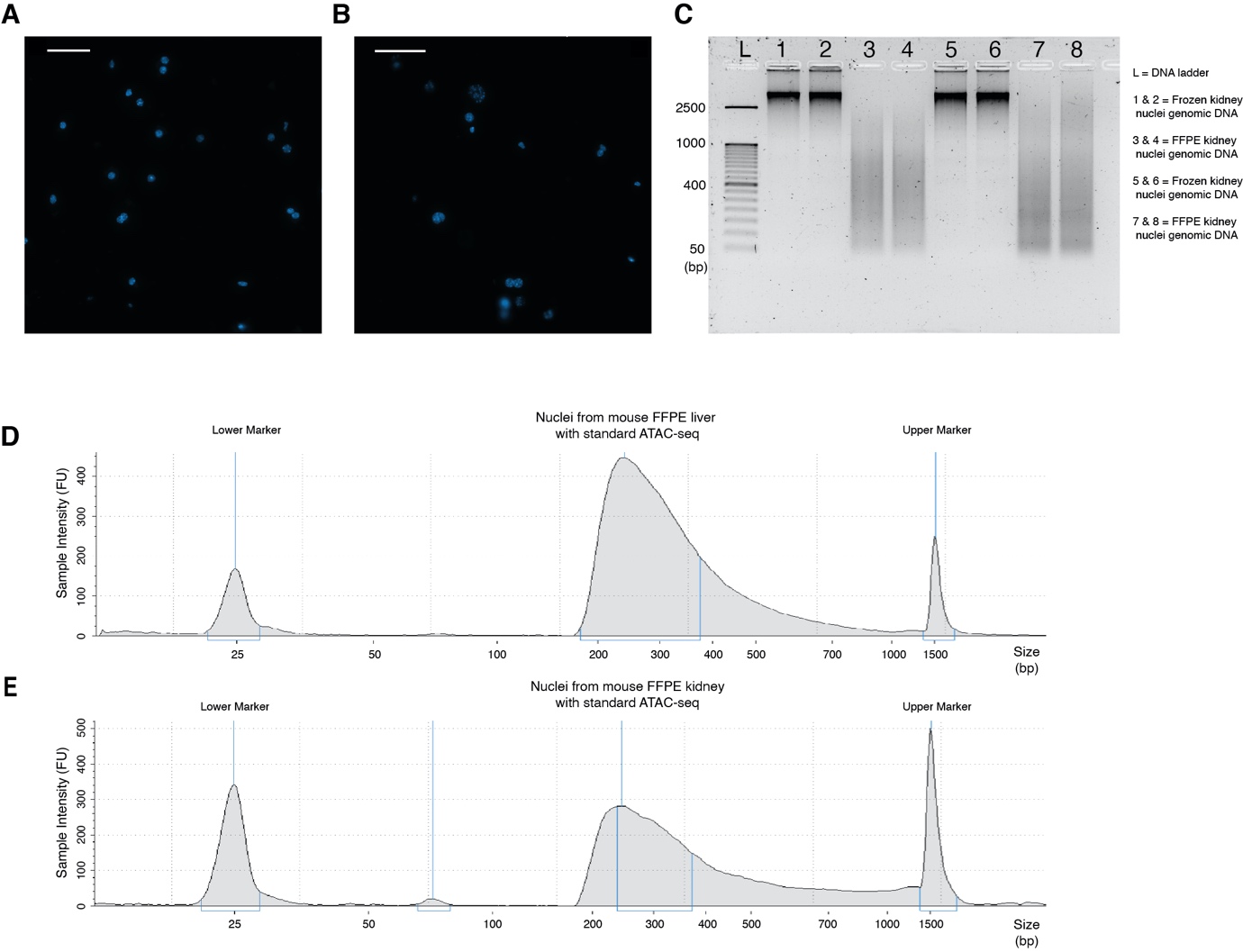

**Supplemental Fig. S1. Standard ATAC-seq on FFPE samples.**

(A) Nuclei purified from 20-µm-thick FFPE mouse liver tissue sections. Scale bar = 50 µm.

(B) Nuclei purified from 20-µm-thick FFPE mouse kidney tissue sections. Scale bar = 50 µm.

(C) Genomic DNA purified from frozen mouse kidney nuclei, frozen mouse liver nuclei, FFPE mouse kidney nuclei and FFPE mouse liver nuclei.

(D, E) Length distribution of PCR amplicons from standard ATAC-seq on FFPE mouse liver nuclei (D) and on FFPE mouse kidney nuclei (E).

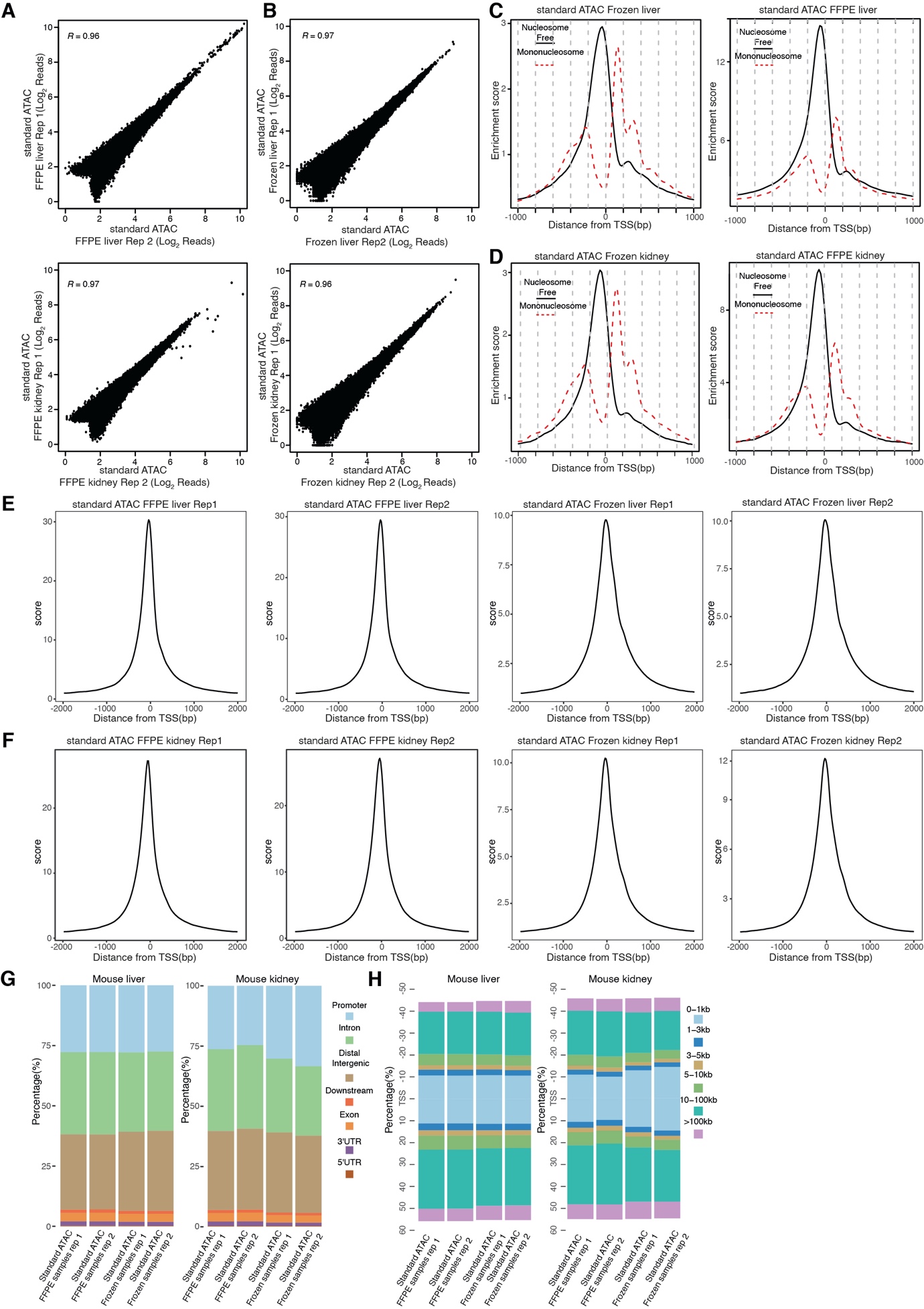

**Supplemental Fig. S2. Comparison of standard ATAC-seq on FFPE samples and standard ATAC-seq on frozen samples.**

(A, B) Genome-wide correlation of two technical replicates in standard ATAC-seq on FFPE mouse liver and FFPE mouse kidney (A), and standard ATAC-seq on frozen mouse liver and frozen mouse kidney (B). *R* = Pearson’s correlation.

(C, D) Sequencing read distribution across transcription start sites (TSS) from nucleosome-free and mononucleosome regions in standard ATAC-seq on frozen mouse samples (left) and standard ATAC-seq on FFPE mouse samples (right).

(E, F) Sequencing read enrichment at transcription start sites (TSS) in standard ATAC-seq on FFPE mouse samples and frozen mouse samples.

(G, H) Genomic annotation of accessible chromatin peaks (G) and distribution of accessible chromatin peaks across transcription start sites (TSS) (H) from standard ATAC-seq on frozen samples and standard ATAC-seq on FFPE samples. Left: mouse liver samples; Right: mouse kidney samples.

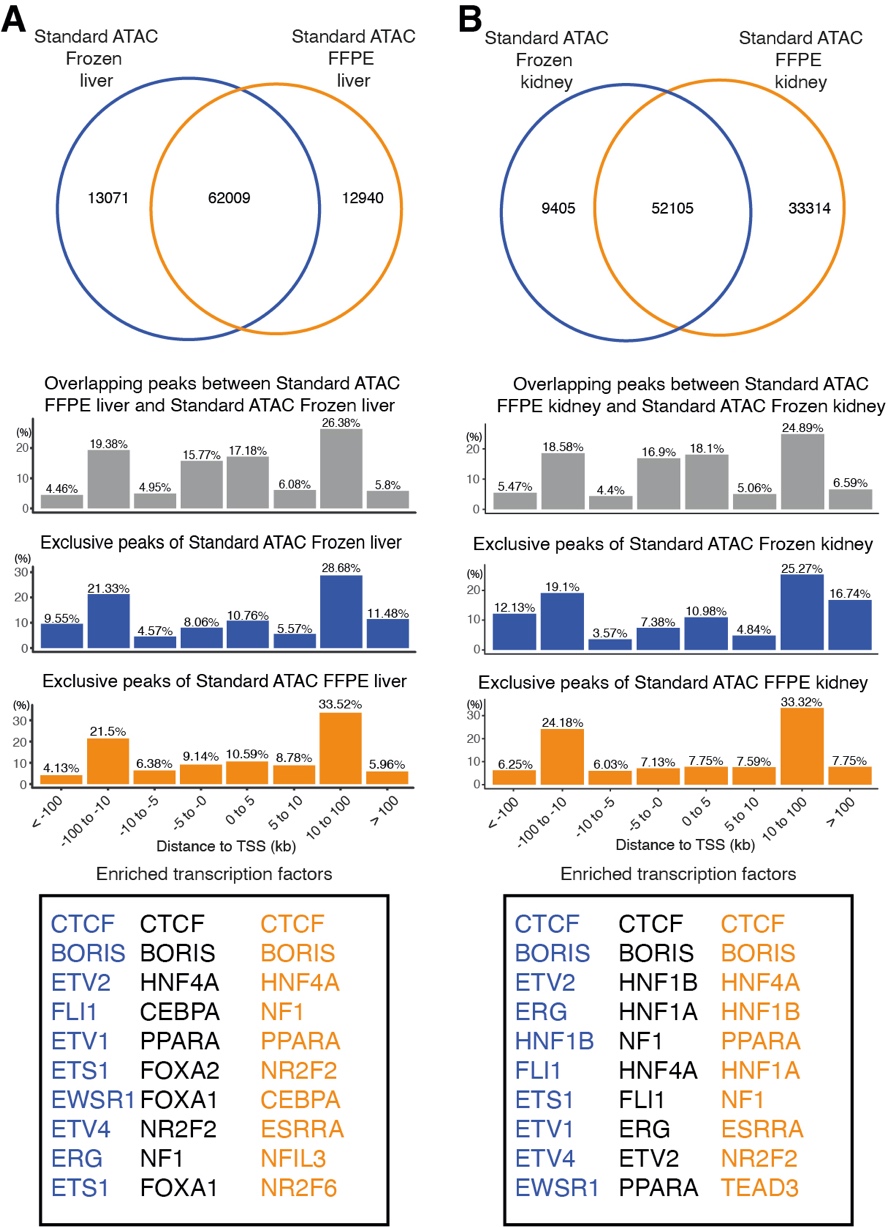

**Supplemental Fig. S3. Comparison of accessible chromatin peaks between standard ATAC-seq on frozen samples and standard ATAC-seq on FFPE samples (** A is for mouse liver, and B is for mouse kidney**).** Top: Overlap of peaks called with the same number of sequencing reads from standard ATAC-seq on frozen mouse sample and standard ATAC-seq on FFPE mouse sample. Middle: Distribution of overlapping peaks and exclusive peaks between standard ATAC-seq on frozen mouse sample and standard ATAC-seq on FFPE mouse sample across transcription start sites (TSS). Bottom: Top 10 transcription factors enriched at overlapping peaks and exclusive peaks between standard ATAC-seq on frozen mouse sample and standard ATAC-seq on FFPE mouse sample.

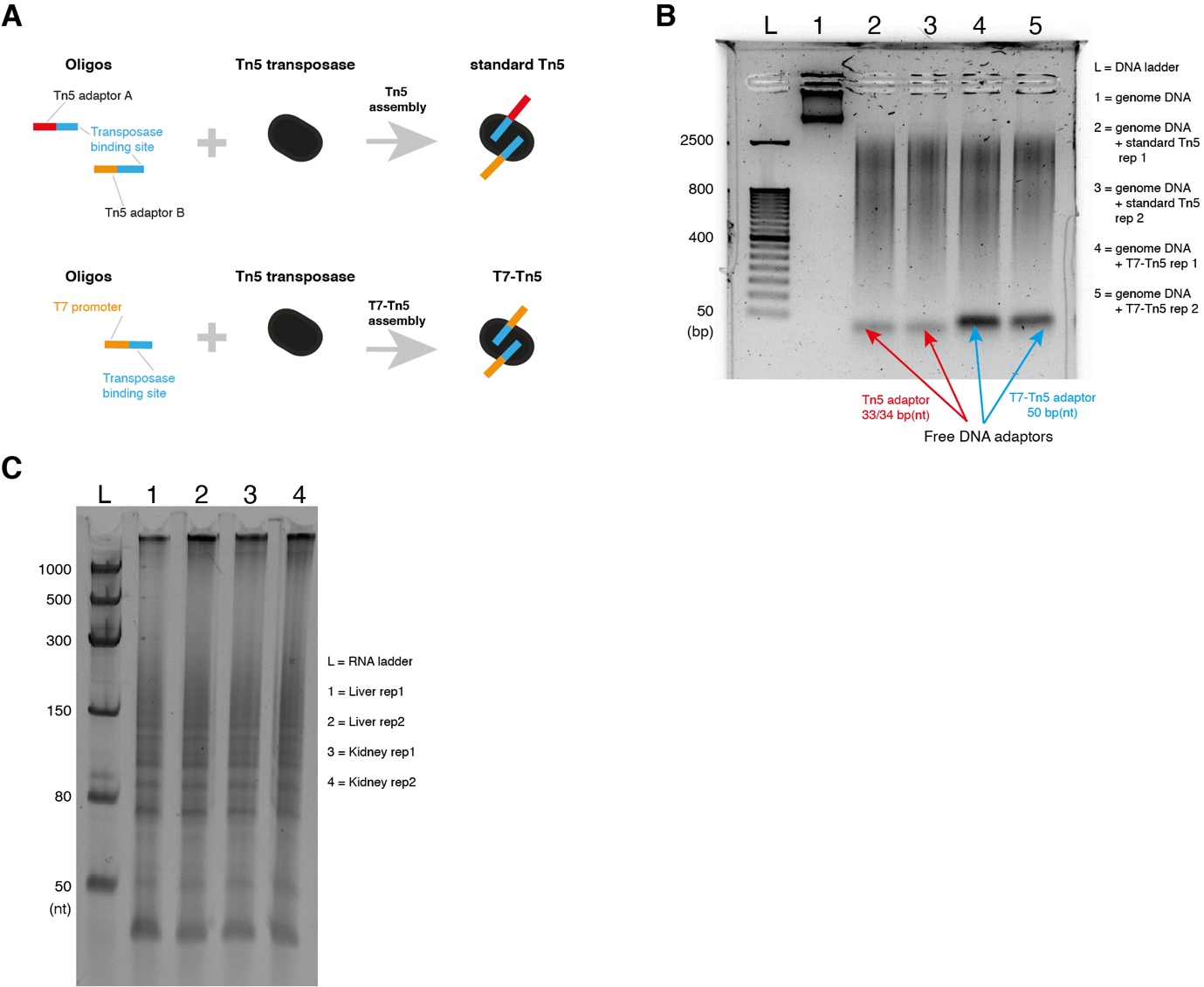

**Supplemental Fig. S4. Combination of T7-Tn5 transposition and T7 *in vitro* transcription in FFPE-ATAC**.

(A) Schematic illustration of Tn5 and T7-Tn5 assembly.

(B) Validation of T7-Tn5 activity. The experiment was repeated three times.

(C) Length distribution of *in vitro* transcribed RNA from mouse FFPE liver and kidney nuclei after T7-Tn5 tagmentation and *in vitro* transcription.

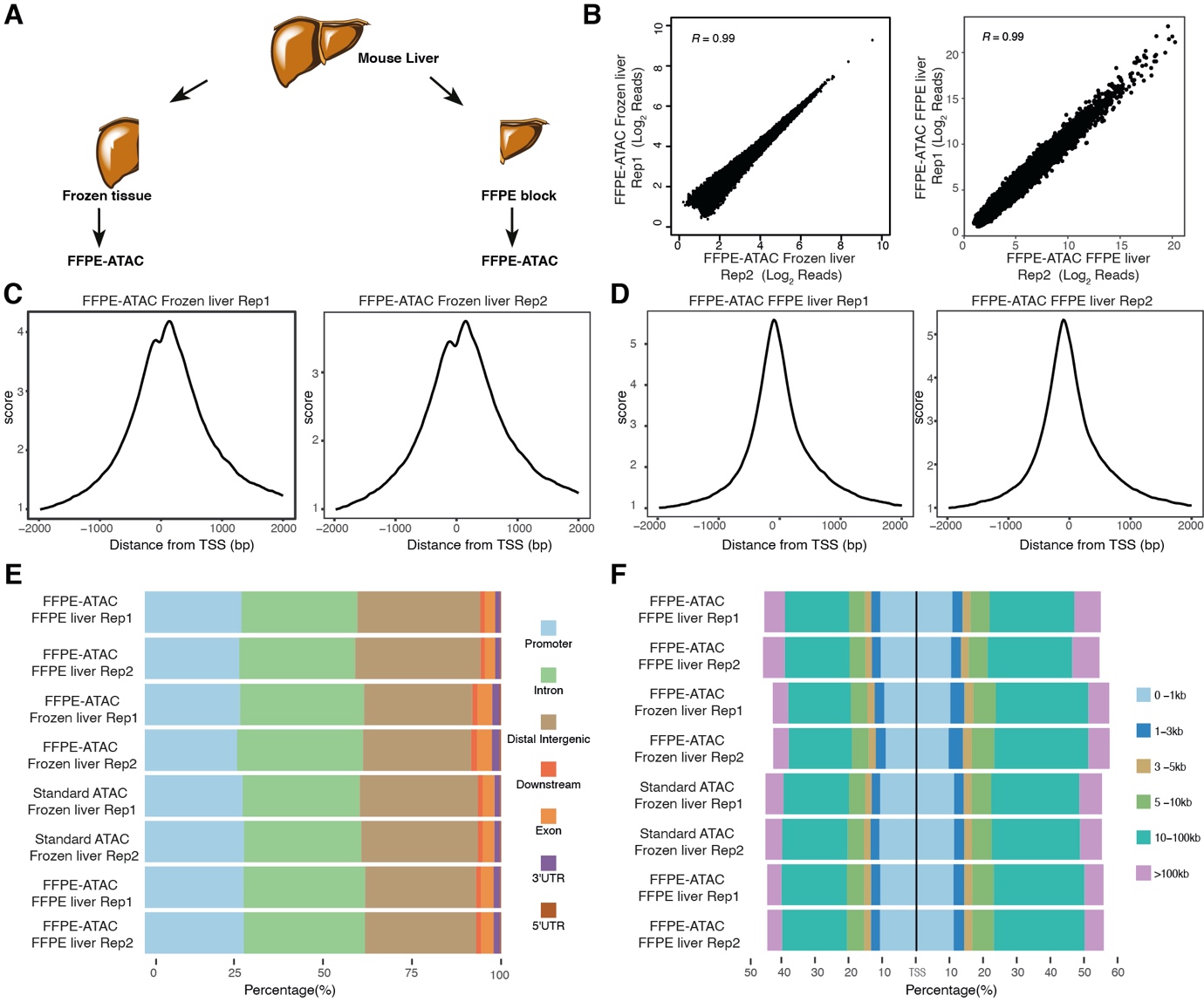

**Supplemental Fig. S5. Comparison of FFPE-ATAC on frozen and FFPE samples from the same mouse liver**.

(A) Experimental design for FFPE-ATAC on frozen and FFPE samples from the same mouse liver.

(B) Genome-wide correlation of two technical replicates in FFPE-ATAC on frozen mouse liver (left) and FFPE-ATAC on FFPE mouse kidney (right). *R* = Pearson’s correlation.

(C, D) Enrichment of sequencing reads at transcription start sites (TSS) in FFPE-ATAC on frozen mouse liver and FFPE-ATAC on FFPE mouse liver.

(E, F) Genomic annotation of accessible chromatin peaks (E) and distribution of accessible chromatin peaks across transcription start sites (TSS) (F) from standard ATAC-seq on frozen mouse liver, standard ATAC-seq on FFPE mouse liver, FFPE-ATAC on frozen mouse liver and FFPE-ATAC on FFPE mouse liver.

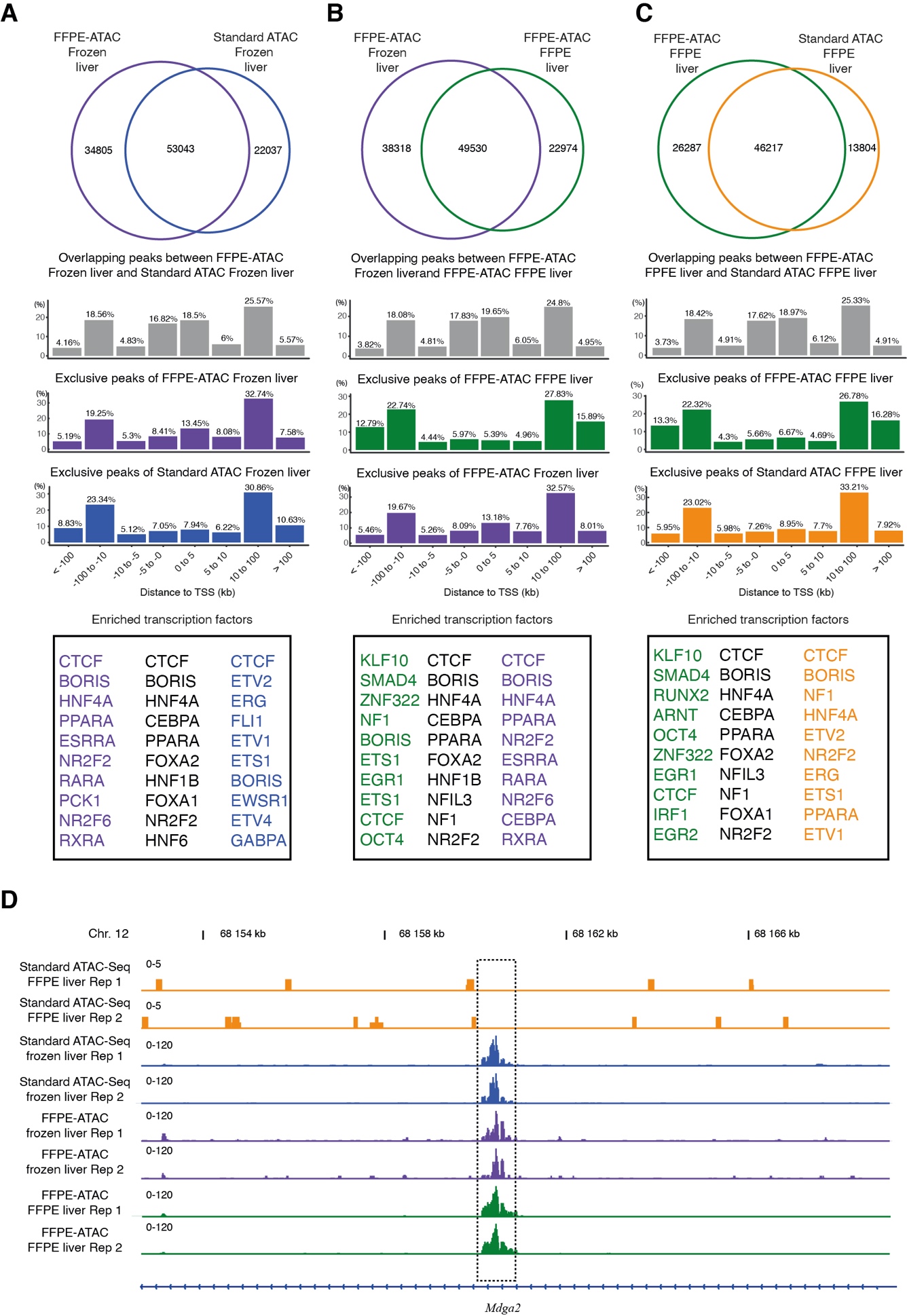

**Supplemental Fig. S6. Comparison of accessible chromatin peaks among different ATAC libraries from frozen and FFPE mouse livers.**

(A-C) Comparison of accessible chromatin peaks among different ATAC libraries:

standard ATAC-seq on frozen mouse liver vs. FFPE-ATAC on frozen mouse liver (A),

FFPE-ATAC on frozen mouse liver vs. FFPE-ATAC on FFPE mouse liver (B),

and standard ATAC-seq on FFPE mouse liver and FFPE-ATAC on FFPE mouse liver (C)**.** Top: Overlap of peaks called with the same number of sequencing reads. Middle: Distribution of overlapping peaks and exclusive peaks from different sequencing libraries across transcription start sites (TSS). Bottom: Top 10 transcription factors enriched at overlapping peaks and exclusive peaks from different sequencing libraries.

(D) A representative peak region, where a accessible chromatin peak were detected from FFPE-ATAC on FFPE mouse liver but no sequencing read counts were amplified by PCR from standard ATAC-seq on FFPE mouse liver. The dotted box indicates the location of differential peak. Gene names are shown at the bottom. Chr. = Chromosome.

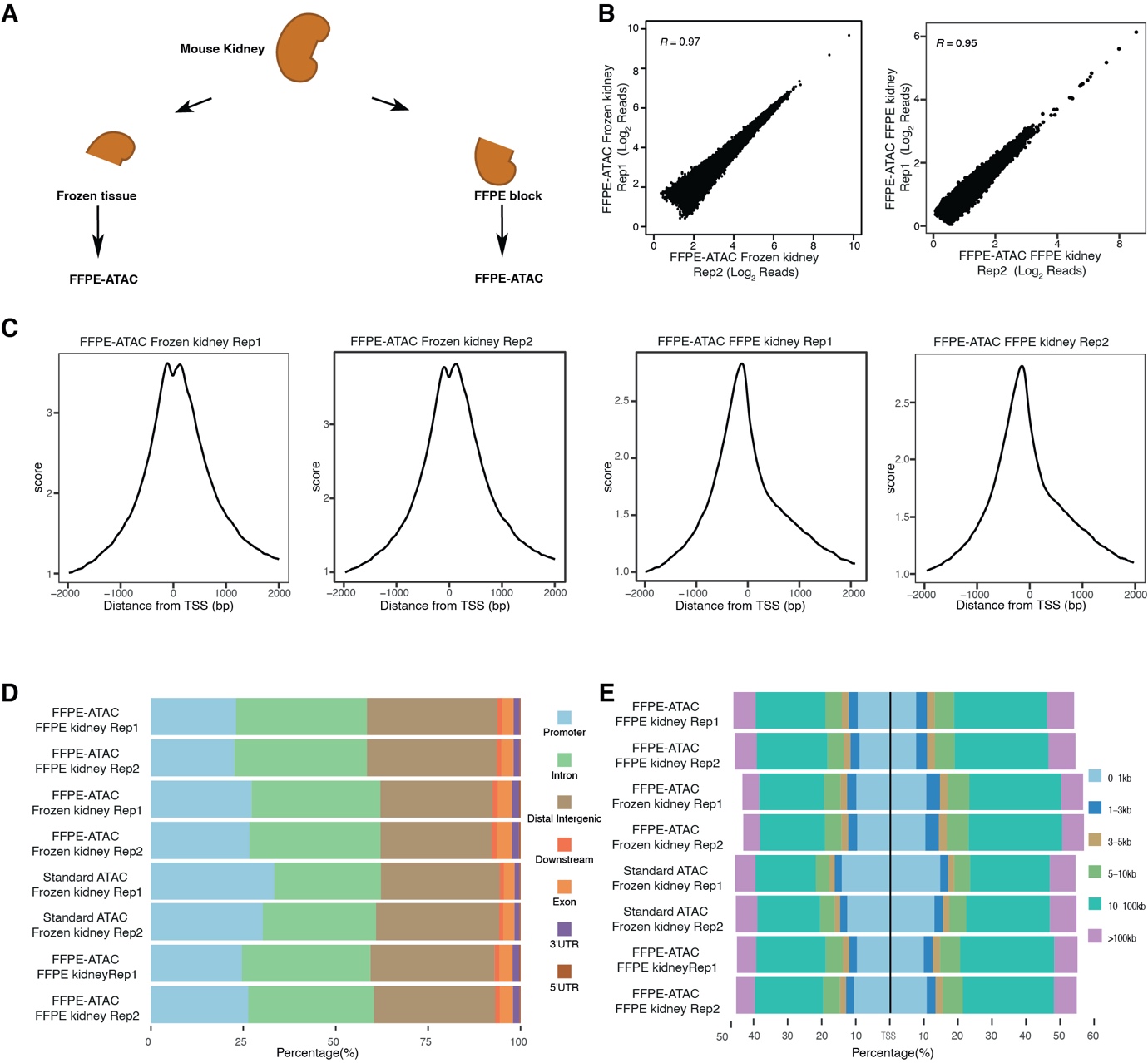

**Supplemental Fig. S7. Comparison of FFPE-ATAC and Omni-ATAC from the same mouse kidney**.

(A) Experimental design for FFPE-ATAC on frozen and FFPE samples from the same mouse kidney.

(B) Genome-wide correlation of two technical replicates of FFPE-ATAC on frozen mouse kidney (left), and FFPE-ATAC on FFPE mouse kidney (right). *R* = Pearson’s correlation.

(C) Enrichment of sequencing reads at transcription start sites (TSS) in FFPE-ATAC on frozen mouse kidney (left two panels), and FFPE-ATAC on FFPE mouse kidney (right two panels).

(D, E) Genomic annotation of accessible chromatin peaks (D) and distribution of accessible chromatin peaks across transcription start sites (TSS) (E) from standard ATAC-seq on frozen mouse kidney, standard ATAC-seq on FFPE mouse kidney, FFPE-ATAC on frozen mouse kidney and FFPE-ATAC on FFPE mouse kidney.

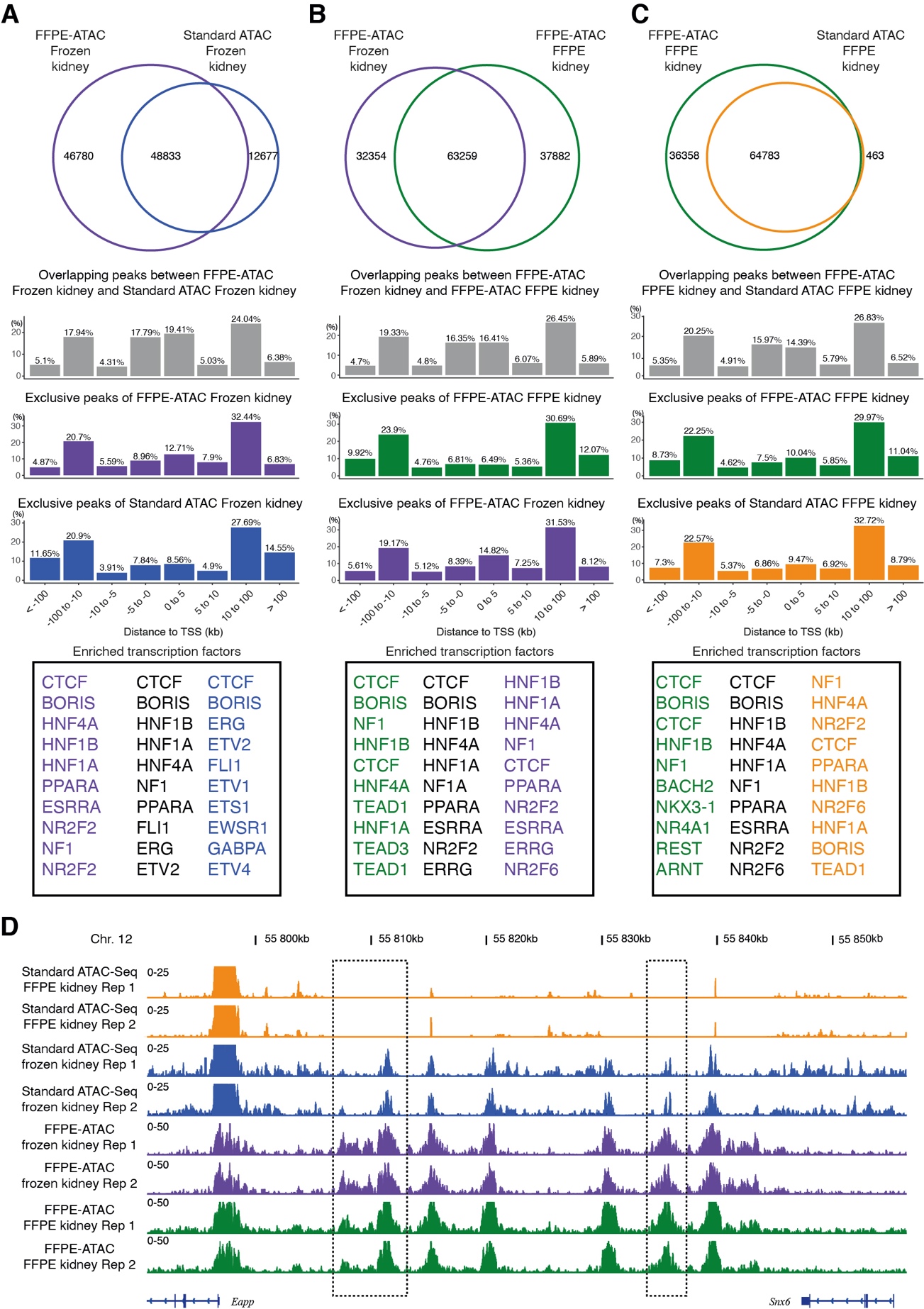

**Supplemental Fig. S8. Comparison of accessible chromatin peaks among different ATAC libraries from frozen and FFPE mouse kidneys.**

(A-C) Comparison of accessible chromatin peaks among different ATAC libraries:

standard ATAC-seq on frozen mouse kidney vs. FFPE-ATAC on frozen mouse kidney (A),

FFPE-ATAC on frozen mouse kidney vs. FFPE-ATAC on FFPE mouse kidney (B),

and standard ATAC-seq on FFPE mouse kidney and FFPE-ATAC on FFPE mouse kidney (C)**.** Top: Overlap of peaks called with the same number of sequencing reads. Middle: Distribution of overlapping peaks and exclusive peaks from different sequencing libraries across transcription start sites (TSS). Bottom: Top 10 transcription factors enriched at overlapping peaks and exclusive peaks from different sequencing libraries.

(D) A representative peak region, where a accessible chromatin peak were detected from FFPE-ATAC on FFPE mouse kidney but no sequencing read counts were amplified by PCR from standard ATAC-seq on FFPE mouse kidney. The dotted box indicates the location of differential peak. Gene names are shown at the bottom. Chr. = Chromosome.

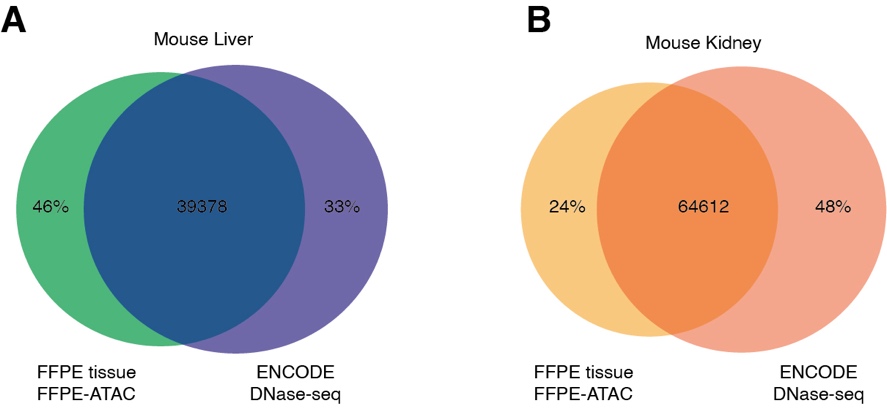

**Supplemental Fig. S9.** Overlap of peaks between FFPE-ATAC on FFPE mouse samples and ENCODE DNase-seq from 8-week-old mouse liver (A) and 8-week-old mouse kidney (B).

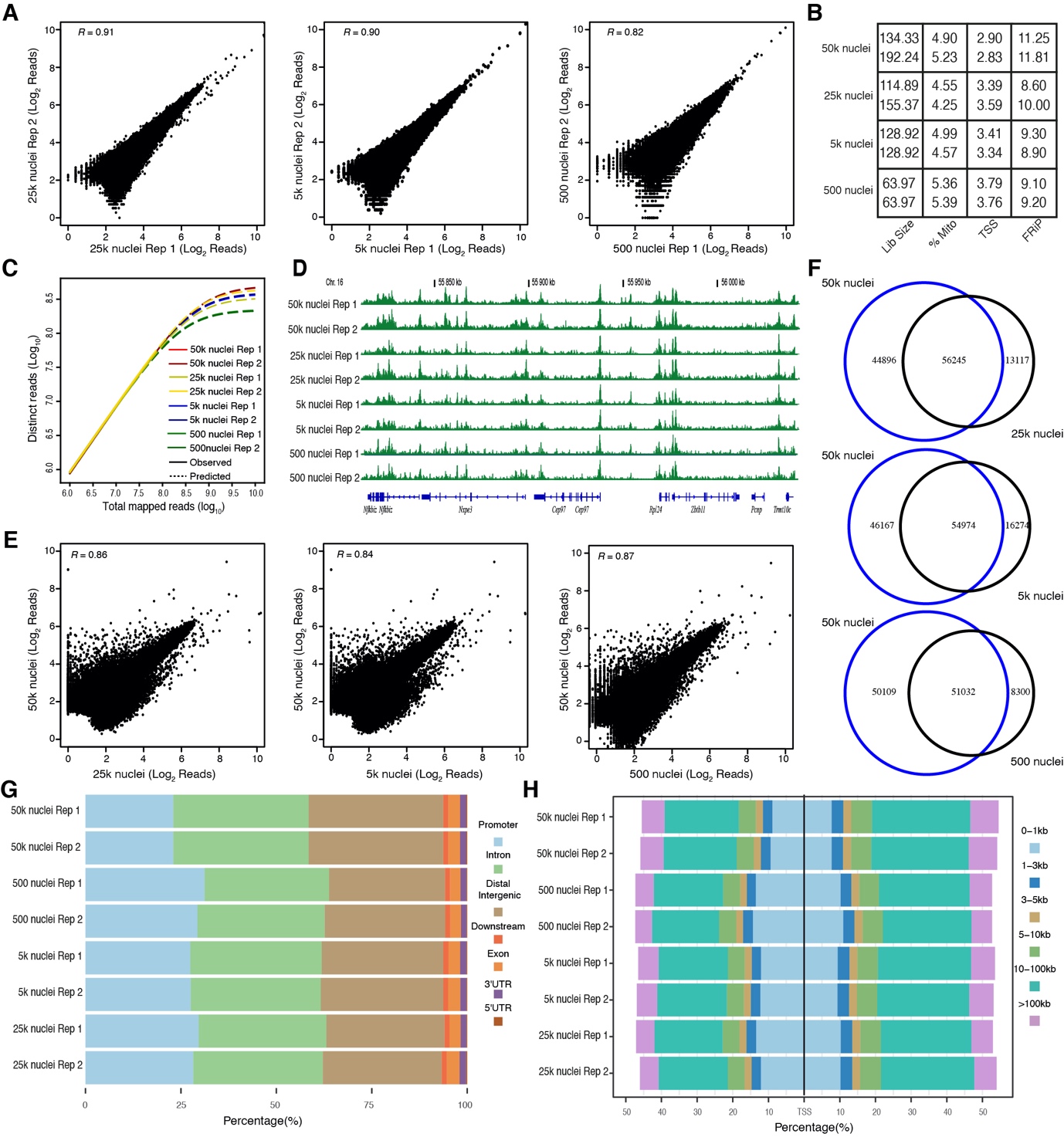

**Supplemental Fig. S10. Sensitivity assay of FFPE-ATAC with nuclei purified from a 20-µm-thick mouse FFPE kidney tissue section**.

(A) Genome-wide correlation of two technical replicates in FFPE-ATAC on FFPE mouse kidney with 25k nuclei (left), 5k nuclei (middle), and 500 nuclei (right). *R* = Pearson’s correlation.

(B-D) Quality control metrics (B), sequencing library complexity (C) and genome browser tracks (D) of FFPE-ATAC on FFPE mouse kidney with different amount of nuclei inputs. Lib size = total sequencing reads of sequencing library (million); %Mito = percentage of mitochondria; TSS = enrichment score at transcription start sites (TSS); FRiP = fraction of reads in peaks. Chr. = Chromosome.

(E) Genome-wide correlation between FFPE-ATAC on FFPE mouse kidney with 50k nuclei and FFPE-ATAC on FFPE mouse kidney with 25k nuclei (Left), 5k nuclei (Middle), and 500 nuclei (Right). *R* = Pearson’s correlation.

(F) Overlap of peaks called with the same number of sequencing reads between FFPE-ATAC on FFPE mouse kidney with 50k nuclei and FFPE-ATAC on FFPE mouse kidney with 25k nuclei (top), 5k nuclei (middle), and 500 nuclei (bottom).

(G, H) Genomic annotation of accessible chromatin peaks (G) and distribution of accessible chromatin peaks across transcription start sites (TSS) (H) from FFPE-ATAC on FFPE samples with different numbers of nuclei.

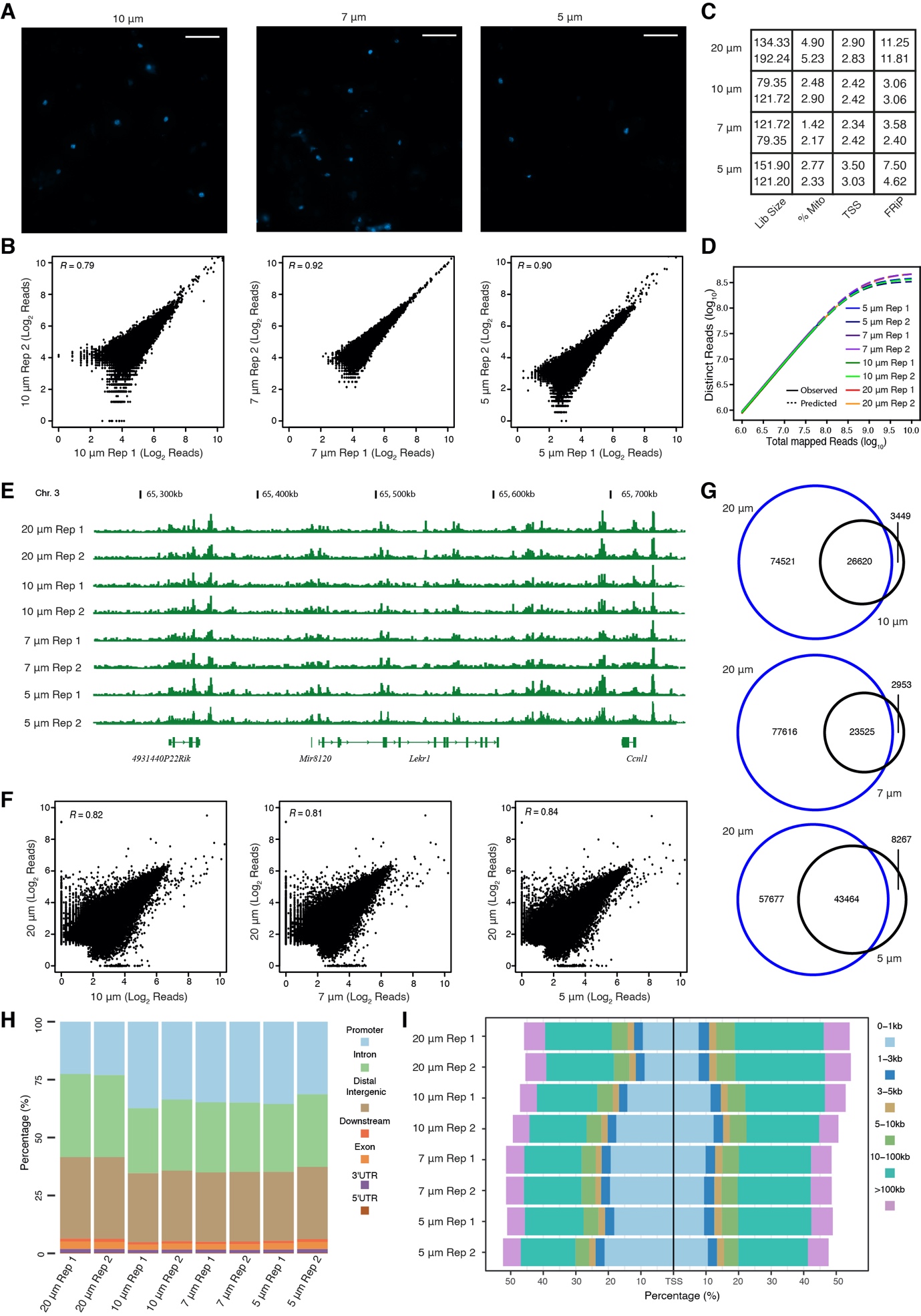

**Supplemental Fig. S11. FFPE-ATAC on FFPE nuclei purified from mouse FFPE kidney tissue sections with different thicknesses**.

(A) Nuclei purified from FFPE mouse kidney tissue sections with thicknesses of 10 µm (left), 7 µm (middle), and 5 µm (right). Scale bar = 50 µm.

(B) Genome-wide correlation of two technical replicates in FFPE-ATAC on FFPE mouse kidney nuclei purified from tissue sections with thicknesses of 10 µm (left), 7 µm (middle), and 5 µm (right). *R* = Pearson’s correlation.

(C-E) Quality control metrics (C), sequencing library complexity (D) and Genome browser tracks (E) of FFPE-ATAC on FFPE mouse kidney nuclei purified from tissue sections with different thicknesses. Lib size = total sequencing reads of sequencing library (million); %Mito = percentage of mitochondria; TSS = enrichment score at transcription start sites (TSSs); FRiP = fraction of reads in peaks. Chr. = Chromosome.

(F) Genome-wide correlation between FFPE-ATAC on FFPE mouse kidney nuclei purified from 20-µm-thick tissue sections and thinner tissue sections: 10 µm (left), 7 µm (middle), and 5 µm (right). *R* = Pearson’s correlation.

(G) Overlap of peaks called with the same number of sequencing reads between FFPE-ATAC on FFPE mouse kidney nuclei purified from 20-µm-thick tissue sections and thinner tissue sections: 10 µm (top, 7 µm (middle), and 5 µm (bottom).

(H, I) Genomic annotation of accessible chromatin peaks (H) and distribution of accessible chromatin peaks across transcription start sites (TSS) (I) from FFPE-ATAC on FFPE nuclei purified from mouse FFPE kidney tissue sections with different thicknesses.

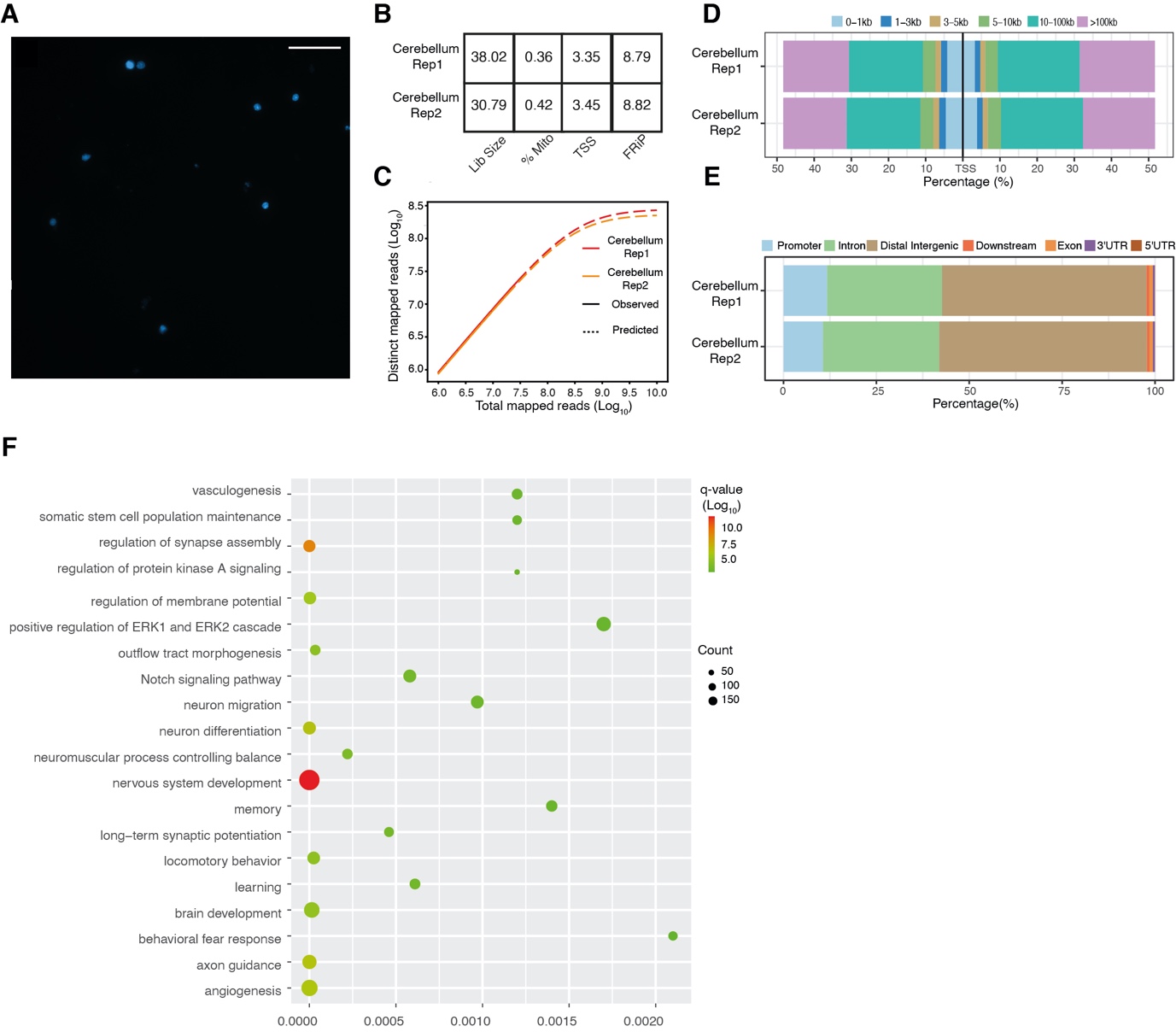
**Supplemental Fig. S12. FFPE-ATAC on nuclei purified from isolated FFPE mouse cerebellum**.

(A) Representative images of nuclei purified from isolated FFPE mouse cerebellum tissue sections. Scale bar = 50 µm.

(B-E) Quality control metrics (B), sequencing library complexity (C), distribution of accessible chromatin peaks across transcription start sites (TSS) (D), and genomic annotation of accessible chromatin peaks (E) from FFPE-ATAC on nuclei purified from the isolated FFPE mouse cerebellum. Lib size = total sequencing reads in the sequencing library (million); %Mito = percentage of mitochondria; TSS = enrichment score at transcription start sites (TSS); FRiP = fraction of reads in peaks.

(F), KEGG pathway analysis of the top 10 000 FFPE-ATAC peaks from the FFPE mouse cerebellum.

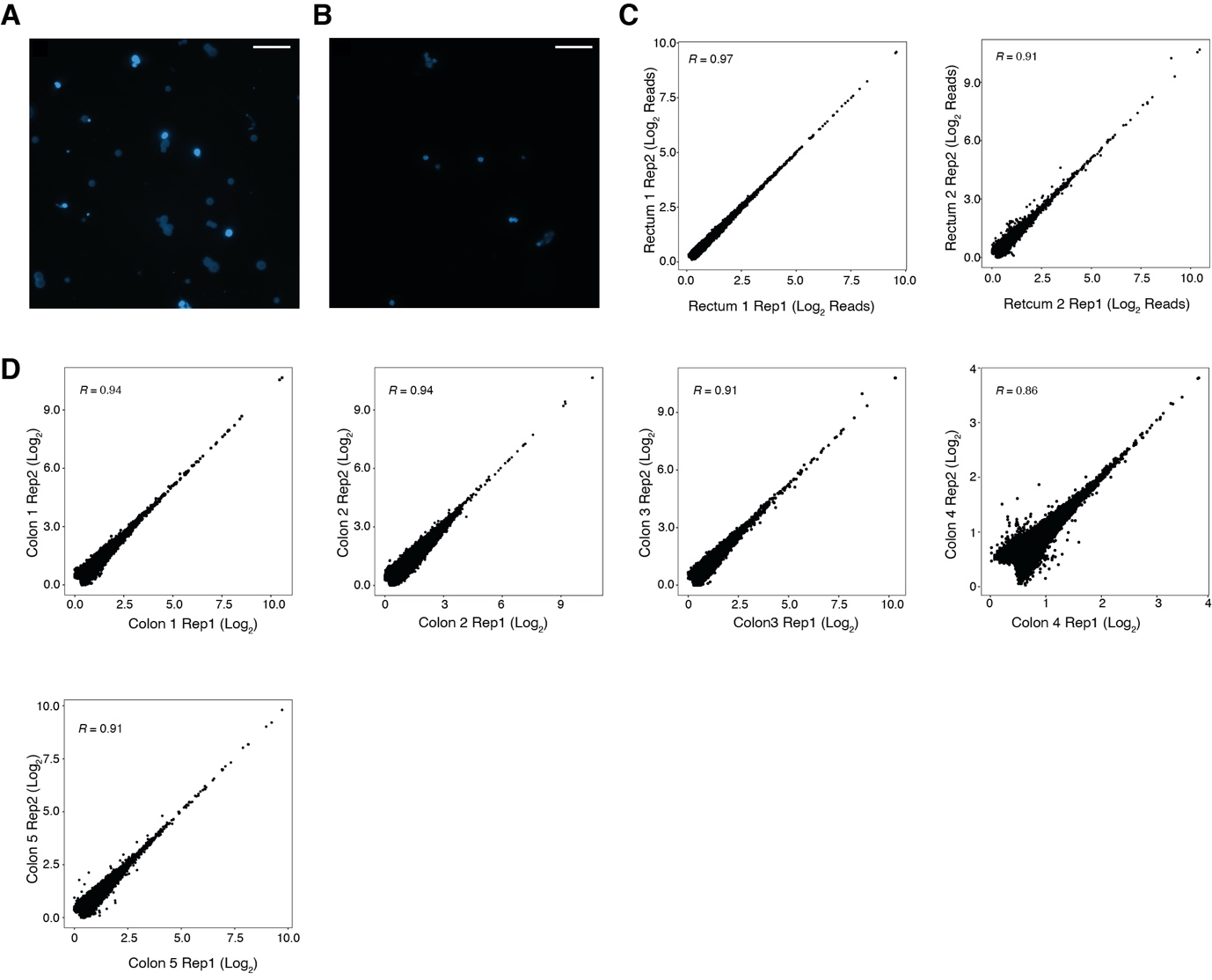

**Supplemental Fig. S13. FFPE-ATAC on nuclei purified from human colorectal cancer (CRC) FFPE tissue sections.**

Representative images of isolated nuclei from FFPE colon cancer (A) and FFPE rectal cancer (B). Reproducibility of FFPE-ATAC from FFPE rectal cancer (C) and FFPE colon cancer (D). Scale bar = 50 µm. *R* = Pearson’s correlation.

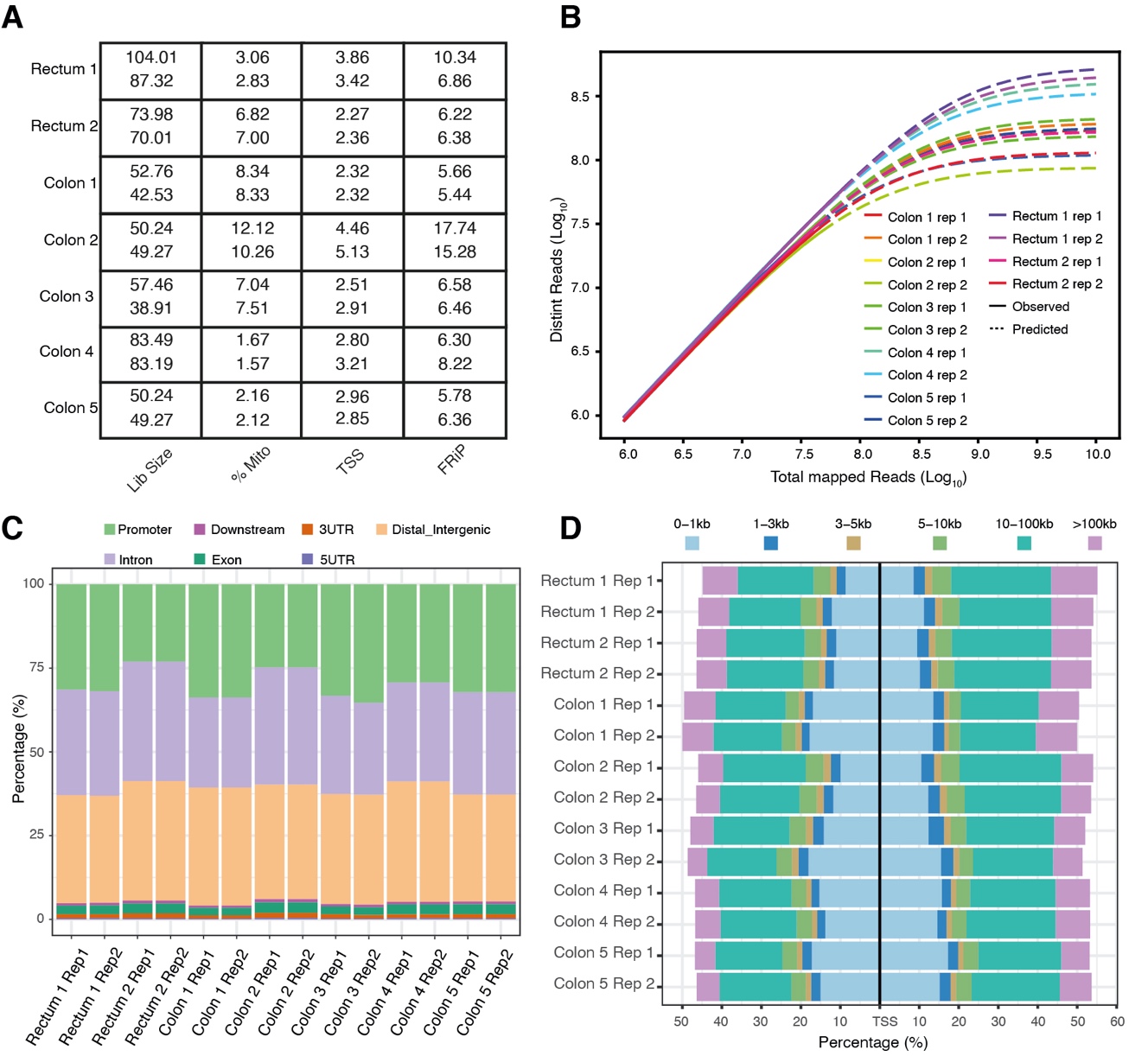
**Supplemental Fig. S14. FFPE-ATAC on 2 cases of FFPE rectal cancer and 5 cases of FFPE colon cancer.**

(A-D) Quality control metrics (A), sequencing library complexity (B), genomic annotation of accessible chromatin peaks (C) and distribution of accessible chromatin peaks across transcription start sites (TSS) (D) from FFPE-ATAC in 2 cases of FFPE rectal cancer and 5 cases of FFPE colon cancer.

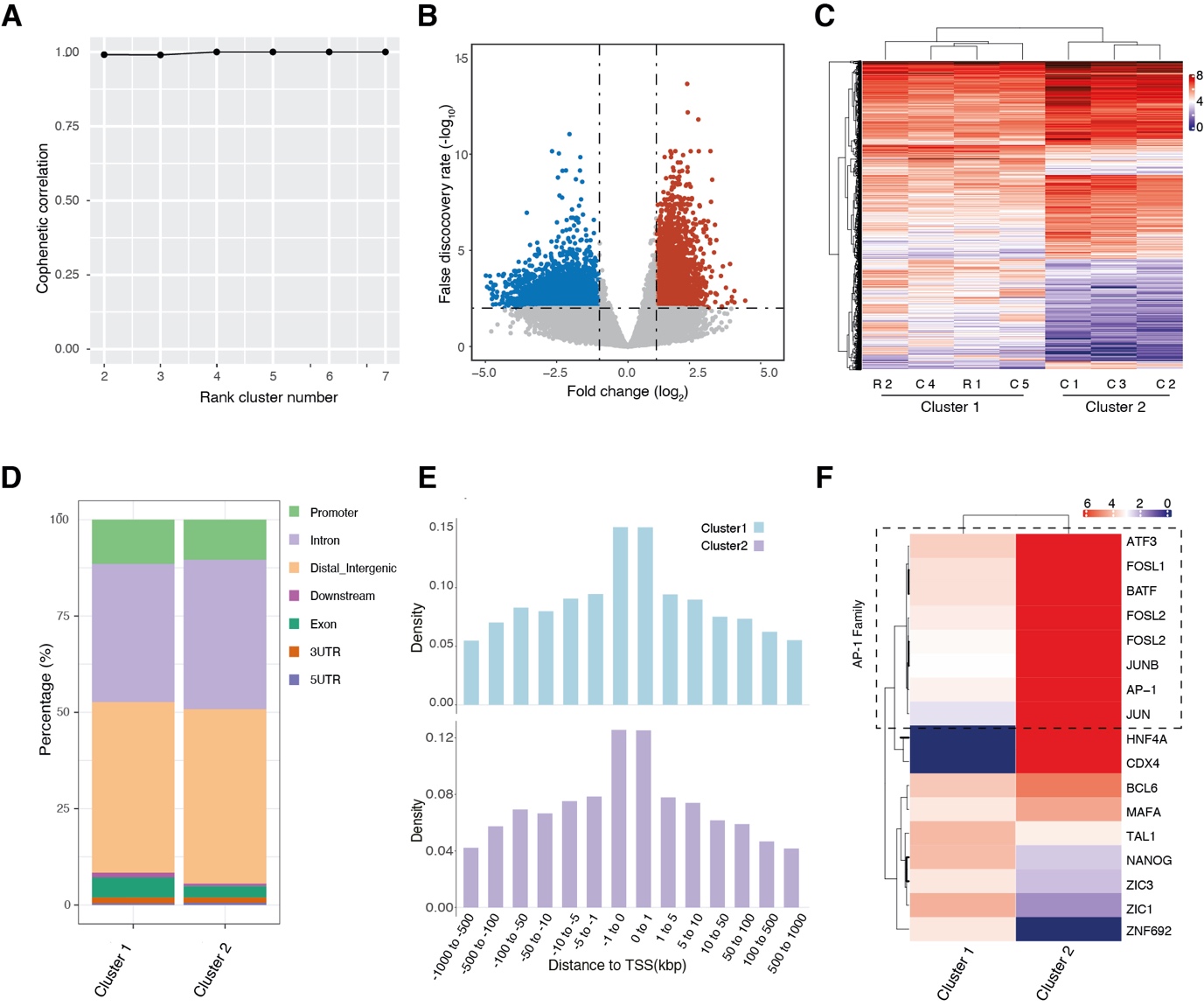

**Supplemental Fig. S15. Decoding chromatin accessible regulatory elements using single clinically archived FFPE tissue sections with FFPE-ATAC.**

(A) Cophenetic correlation coefficient corresponding to the consensus matrices for cluster numbers from 2 to 7, where we chose 2 as the cluster number.

(B) Volcano plot showing differential FFPE-ATAC accessible regions between two clusters, where each dot represents one peak region. The significantly differential peaks are color-coded by the parameters of fold change > 2 and false discovery rate < 0.01. Blue = cluster 1-specific; red = cluster 2-specific.

(C) Heatmap showing the clustering of significantly differential FFPE-ATAC peaks among the two clusters.

(D) Genomic features of the significant FFPE-ATAC peaks from the two clusters, where we did not find significantly different features.

(E) Density histograms representing the distribution of significantly differential FFPE-ATAC peaks among the two clusters.

(F) Cluster of the top 10 ranking significantly enriched transcription factors from the two clusters.
